# Interactive auditory task reveals complex sensory-action integration in mouse primary auditory cortex

**DOI:** 10.1101/2022.12.12.520155

**Authors:** Ji Liu, Patrick O. Kanold

**Author notes:** Corresponding author: Patrick O. Kanold, Dept. of Biomedical Engineering, Johns Hopkins University, 379 Miller Research Bldg., Baltimore, MD 21205 USA.

## Abstract

Predictive coding theory postulates that the brain achieves perception by actively making predictions about the incoming sensory information and correcting them if errors signals arise. These signals are likely the most relevant when the individual is actively interacting with the environment and where the sensory outcome determines the ongoing action. In addition, the cerebral cortex is thought to play a key role in generating these signals. Thus, to study the representation of error signals in the primary sensory cortex, we trained mice to perform an interactive auditory task that coupled their actions to the generated sound and perturbed this coupling to evoke putative error responses. We imaged Layer 2/3 (L2/3) and Layer 4 (L4) neurons in the mouse primary auditory cortex, and we identified not only neurons that mainly encoded action related information but also neurons encoding the mismatch between the action and the sound. These results show that a subset of A1 neurons encode the nonlinear interactions between the sound and the action. Furthermore, more L2/3 neurons encoded action related information than L4, indicating that action-sound integration emerges hierarchically in A1 circuits. Together, our results show that complex interactions between action and sound happen in A1 and that some A1 neurons responses reflect the violation of the learnt relationship between the action and sound feedback. Thus, primary sensory cortices not only encode sensory driven activity but also represent the complex interplay between sensory inputs, expectations, and errors.

## Introduction

Sensory perception is a complex process that requires active involvement and interaction of several brain areas. The predictive coding theory is a framework for explaining perception, which postulates that the brain perceives the outside world by actively making predictions according to its own model of the world, and constantly updates the model via interactions between lower and higher order brain regions through error signals (Friston, 2010; Heilbron and Chait, 2017) (Figure 1A). Thus, a model of the world is learned and used to make predictions on the effects of the individual’s interaction with the world. For example, if motor actions are evoking sounds, e.g., playing a piano, then the desire to evoke a particular tone (desired sound D in Figure 1A) leads to a specific motor plan (M) and the corresponding motor action, which constitutes the inverse model for motor control (Kawato, 1999) (Figure 1A). Due to the slowness of the sensory feedback, it is unstable to solely use sensory information for feedback control. Rather, the forward model that predicts the sensory outcome of the planned action is needed. In the piano playing example, the motor plan to generate the desired sound D would be fed into the forward model to generate the predicted sound P. The evoked sound (S) is then compared with the predicted sound P. If the evoked sound does not match the prediction an error signal (E) is generated leading to an adjustment of the predictions. However, where predictions and error signals are computed in the brain has been debated. Importantly, error signals most likely arise when individual organisms actively interact with their environment to acquire new information to update the brain’s prediction of the outside world. Therefore, testing predictive coding theory and identifying the underlying neural substrate requires experimental paradigms that explicitly incorporate actions into sensory processes.

**Figure 1.**
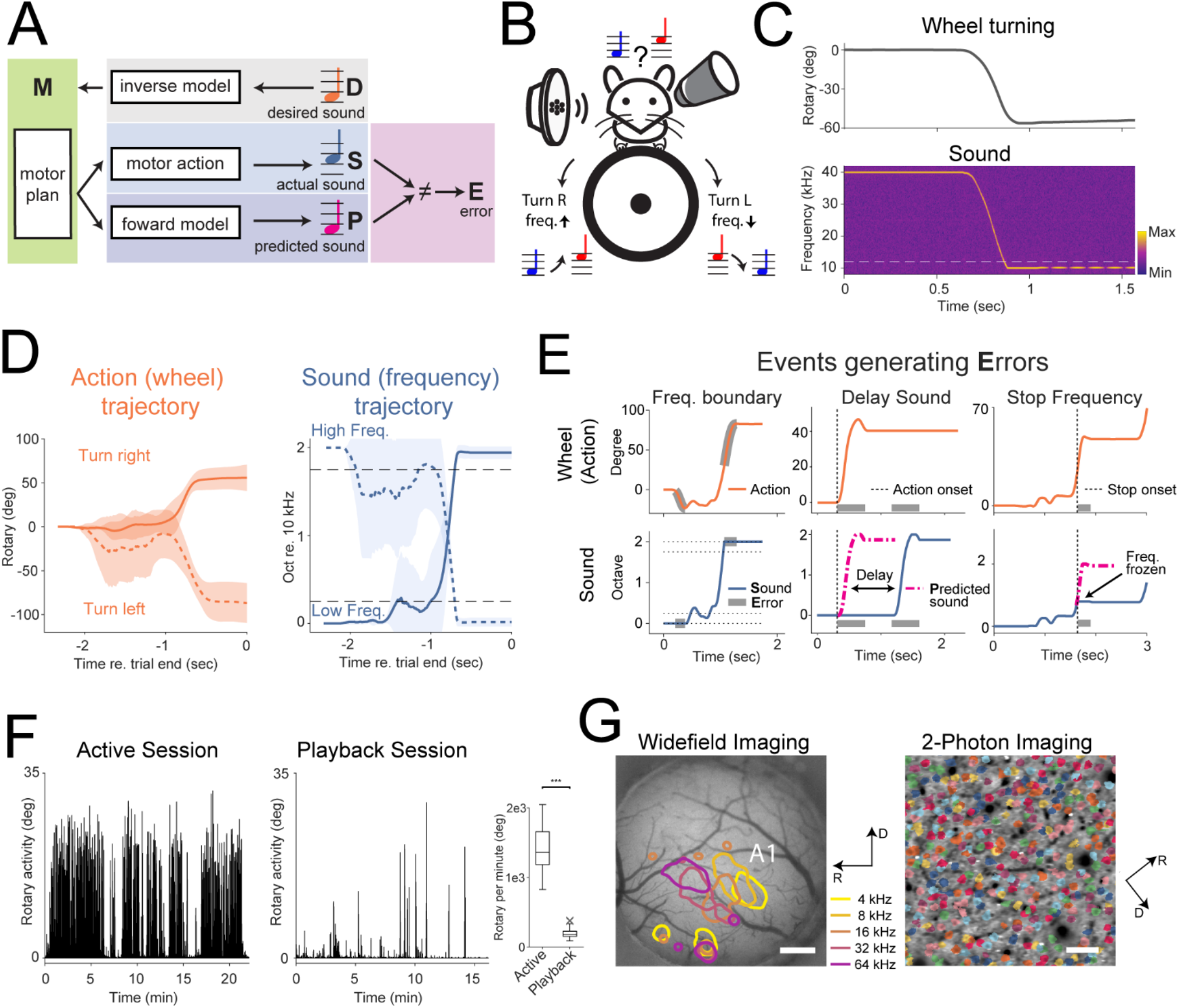
Mice were trained to perform an interactive behavior paradigm. (A) The internal model of motor planning suggests the existence of both the inverse model, which generates the motor plan according to the desired sensory outcome (designed sound D), and the forward model, which generates a prediction of the sensory outcome (predicted sound P) based on the motor plan. The error signal (E) arises when the actual sound (S) fails to match the predicted sound P. (B) Schematic of the behavior paradigm. The mouse was trained to modulate the carrier frequency by turning the wheel. (C) Example action and frequency stream from one hit trial. Top: the rotary reading corresponding to the left turning action. Bottom: the microphone-recorded spectrogram of the frequency stream. The white dash line indicates the boundary of the target zone for this trial. After the hit condition was met, the carrier frequency was frozen for 0.5 second while a 10 Hz amplitude modulation was added to signal the trial outcome. (D) Average wheel and frequency trajectories of hit trials with short hit latencies (⩽ 2 seconds) from one mouse in one active session. The solid lines correspond to trials with a low start frequency while the dashed line correspond to trials with a high start frequency. The solid and dashed lines represent the mean and the shaded regions represent standard deviation. (E) Three forms of sound-action mismatch were studied. All three forms involved the decoupling between wheel turning and the carrier frequency change. Left: errors happened when the frequency streams were clipped by the frequency lower and upper boundaries, which was intrinsic to the paradigm design. Middle and right: two forms of random perturbations were artificially introduced. Middle: the Delay-Sound (DS) perturbation introduced a delay (1 second) between the wheel turning and the corresponding carrier frequency change. Right: the Stop-Frequency (SF) perturbation froze the frequency stream briefly at one of the predetermined frequencies. The vertical dotted line indicates the onset of the SF perturbation. The actual frequency stream was held constant for 0.25 second and deviated from the predicted frequency stream (violet dot-dash line). The gray bars indicate the period these errors occurred. (F) Left and middle: example wheel movements from one active session and the subsequent playback session. Right: mice showed significantly less movement in playback sessions than in active sessions (Wilcoxon rank sum test, p=3.3×10^−18^). (G) Left: contour plots showing “hotspot” of fluorescence increase evoked by different tone frequencies during widefield imaging. A1 was identified through its caudolateral-mediodorsal tonotopic gradient. The scale bar represents 500 μm. Right: A1 neurons under 2P microscopy. The example image shows a cropped region from the full FOV for clarity. Neurons were colored according to segmentation by Suite2P. The scale bar represents 50 μm. The rotation of the 2P FOV was due to the rotation of the nosepiece relative to the scanning directions.

Studies in the visual system have shown that action influences sensory processing as locomotion increases visual responses in the mouse visual cortex (Mineault et al., 2016; Niell and Stryker, 2010; Pakan et al., 2016). Furthermore, error signals were evoked in mouse primary visual cortex by the mismatch between expected visual flow and actual feedback (Keller et al., 2012), lending evidence for the predictive coding theory. In the auditory system, this framework has been proposed to explain phenomena such as Stimulus Specific Adaptation (SSA) and Mismatch Negativity (MMN) (Carbajal and Malmierca, 2018), where less common oddball stimulus evokes larger neuronal responses along the auditory pathway than the common standard stimulus (Parras et al., 2017). These responses have been largely interpreted as error signals for violating auditory expectations. However, these paradigms typically present auditory stimuli in a passive setting and are thus not interactive in nature and do not require the active involvement of the subject. In contrast, a closed-loop experimental design, where the sensory outcome depends on the action of the animal, might better engage the predictive processing. Although closed-loop designs have been used in studying auditory perception (Nelson et al., 2013; Rummell et al., 2016; Schneider et al., 2018), these studies employed simple relationships between the action and the sound and did not tightly couple action and sensory feedback. Since action modulation in sensory cortex depends on experience (Attinger et al., 2017), by using a tighter action-sensory coupling we could potentially identify the complex interactions between the two that give rise to error signals in ACX. Furthermore, ACX neurons are heterogeneous (Bandyopadhyay et al., 2010; Rothschild et al., 2010; Winkowski and Kanold, 2013) and encode information beyond the properties of the sound stimulus such as behavioral choices and reward (Brosch et al., 2011; Francis et al., 2018; Guo et al., 2019). We thus postulate that ACX also encode error signals.

Thus, to test if ACX neurons encode error signals we designed a novel behavioral paradigm that allowed mice to directly interact with the presented sound. Specifically, we trained mice to turn a wheel, whose readout was coupled to the increase or decrease of the carrier frequency of the sound stream the mice were presented with (Figure 1B, C). Mice needed to “steer” the carrier frequency either into the low or the high end of the sound spectrum (target zones) for water reward (Figure 1D). This paradigm allowed mice to continuously evaluate and control one attribute of the sound, i.e., frequency. We studied the three error-inducing scenarios where the sound frequency mismatched with the animal’s action (Figure 1E) which included two forms of perturbations: Delay-Sound (DS) and Stop-Frequency (SF) where we disrupted the default relationship between the action and the sound by delaying the sensory outcome or freezing the carrier frequency (Figure 1E). To study neural responses in this paradigm, we used 2-Photon (2P) imaging of the primary ACX (A1) to investigate whether potential error signals evoked by the perturbations were present. Because the cortex performs hierarchical computations (Winkowski and Kanold, 2013), we imaged both Layer 4 (L4) and Layer 2/3 (L2/3) to investigate if the error signal emerged in specific layers. We identified groups of neurons in L2/3 and L4 of A1 that were responsive to distinct features including sound or action with L2/3 containing more neurons responsive to actions than L4, suggesting the presence of a forward model in A1 which potentially encodes the predicted sensory outcome from the action. Further, we found a group of neurons that encoded error signals by responding to the decoupling (DCP) between action and sound. Our results suggest that A1 represents the complex relationship between action and sound in our paradigm as DCP responses signal the violation of the default rules. Thus, error signals for sensory predictions are present in primary sensory cortices.

## Results

### Mice were trained to perform an interactive auditory task

To investigate how A1 neurons encode the interaction between action and sensory feedback and how such interactions emerge across the cortical layers, we first trained mice to perform an interactive auditory task (n=6 mice, 3 male and 3 female). Our interactive task required the animal to change the frequency of a sound into a target frequency range. The mice were trained to control the frequency of the sound by turning a wheel placed beneath their front paw. The mice were presented with a frequency modulated sound stream and its carrier frequency would decrease or increase as the animal turned the wheel clockwise or counterclockwise (Figure 1B). The sound stream started either at the low end (10 kHz) or at the high end (40 kHz) of the predefined spectral range. The task required the animal to “steer” the low starting frequency (10 kHz) into the high target zone (33.6 kHz to 40 kHz, 0.25 octave in width), or “steer” the high starting frequency (40 kHz) into the low frequency target zone (10 kHz to 11.9 kHz, 0.25 octave in width). The carrier frequency would not decrease or increase beyond the low and high frequency boundary (10 and 40 kHz, respectively). Figure 1C shows the spectrogram of the microphone-recorded sound stimulus of one trial with the high starting frequency, along with the action of the animal as shown by the rotary reading. Figure 1D shows the average frequency and rotary trajectories for hit trials with short hit latencies (within 2 seconds). These results show that the animals turned the wheel robustly to perform the task.

Immediately following each active session, we presented animals with a playback session. In playback sessions, we first played pure tones to measure tuning curves of imaged neurons, following which we presented the animal with a subset of the frequency stream generated in the active session as a control condition. The playback session allowed us to measure the contribution of stimulus selectivity to neural responses with minimum confound from animal movement as previous studies suggest that auditory responses could suppressed by the animals’ action (Rummell *et al*., 2016; Schneider *et al*., 2018). In playback sessions, the animals remained relatively still and showed much less movement (Figure 1F). The animals were able to perform the task well during imaging sessions. The overall hit rate of all behavioral sessions with simultaneous imaging was 70.2% ± 8.9% (Mean ± STD). The maximum hit rate over any consecutive 100 trials during one session was 75.6% ± 10.0%. The average time to achieve a hit was 1.05 second ± 1.03 second.

### A1 neurons show both sensory and action driven activity during an interactive auditory task

We identified A1 with widefield imaging and performed 2P imaging of neural activity in A1 L2/3 and L4 (Figure 1G; L2/3, 26 field of views (FOVs), 46328 neurons imaged; L4, 25 FOVs, 53211 neurons imaged). Given the interactive nature of the task, neuronal activities likely contain contributions from both sensory and action signals. Thus, to understand how neural responses encode action and the corresponding sensory outcome, we constructed a linear model whose predictors included energy in different frequency bins (F), the rate of frequency modulation (FM) and wheel turning (W, +W and -W for right and left turns respectively, Figure 2A). As we hypothesize that neurons could show sensitivity to the synergy between W and F we added an interaction term between the two (Figure 2A, see Methods). To account for the behavioral state dependence of the neuronal responses, we introduced one task term (T) that took value of 1 for predictor values constructed from active sessions and −1 for predictor values constructed from playback sessions. The T term alone can account for a shift of baseline activity levels across the behavioral states, while the interaction terms between T and sound encoding terms, i.e., F and FM, can account for sound driven responses that are behavioral state dependent (FT and FMT term). Finally, we added one term that accounts for the reward consumption (Figure 2A, Rw). Thus, this model has 8 groups of predictors that encode various features during the active and playback sessions (Figure 2B).

**Figure 2.**
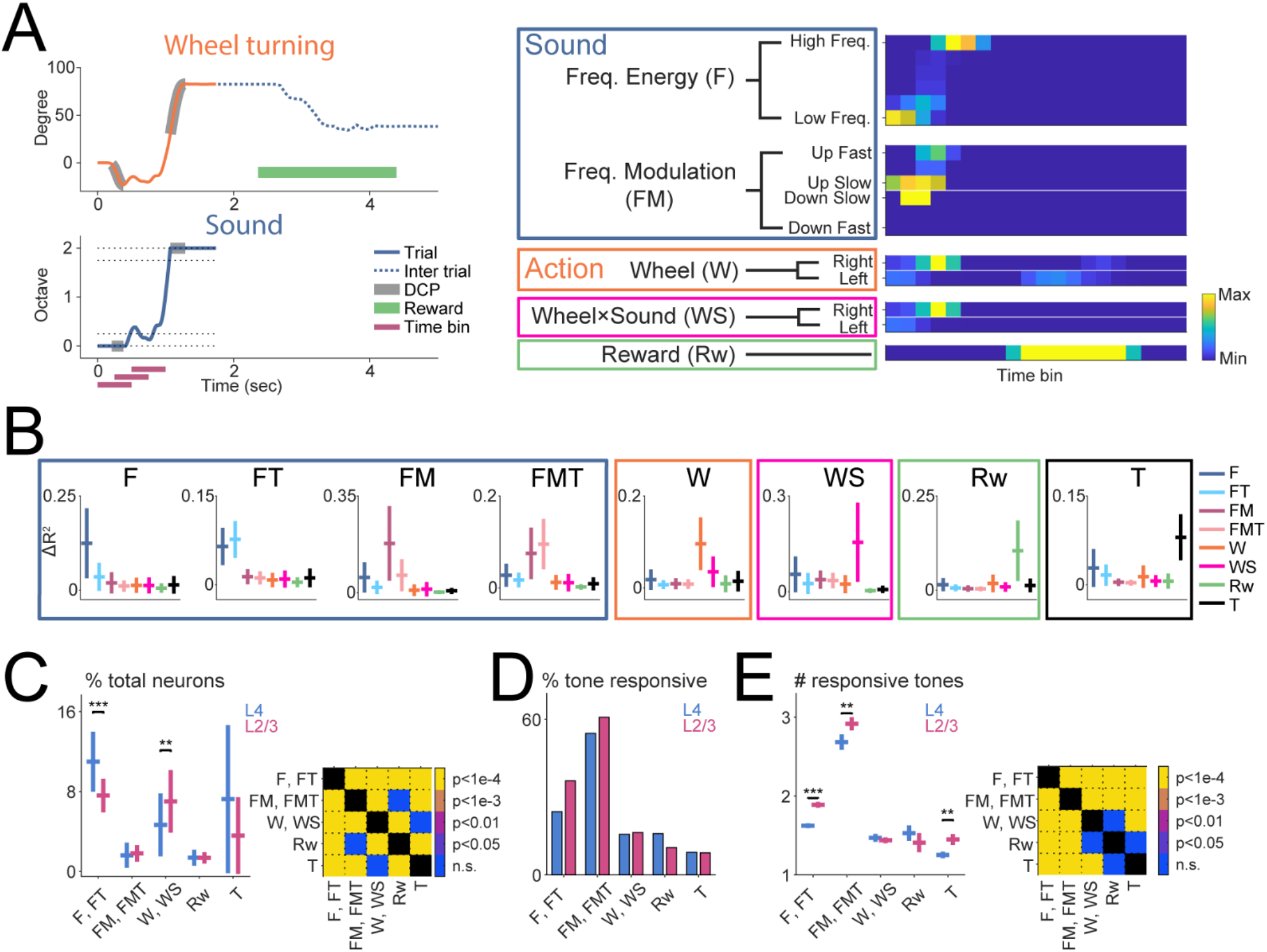
Multiple non-sensory factors contribute to A1 responses. (A) Left: wheel trajectory (top) and frequency trajectory (bottom) of an example hit trial. The part of action that was not translated into carrier frequency changes was marked in black (decoupling, DCP). The green line marks the post-trial period when the animal consumes the water reward. The wheel movements were recorded both in trial (top, solid blue curve) and inter-trial (blue dotted curve). The purple lines represent the time windows (0.5 second in duration) from which we constructed the predictors for the linear model. Each window was shifted by 0.25 second from the previous one. Right: the predictors constructed from the same trial shown on the left. (B) The group of neurons sensitive to various features were identified through their maximum group ΔR^2^ term. (C) The faction of feature sensitive neurons as a function of cortical depth. L4 vs L2/3 Wilcoxon rank sum test (significance adjusted with Bonferroni correction): F, FT, p=1.8×10^−5^; FM, FMT, p=0.22; W, p=0.0086; WS, p=1.8×10^−4^; Rw, p=0.55; T, p=0.015. (D) The fraction of neurons within each feature group that were tone responsive as a function of layer. (E) Left: within the tone responsive neurons of each feature group, the number of tones evoking significant responses in each neuron were plotted as a function of layer. L4 vs L2/3 Wilcoxon rank sum test: F, FT, p=1.2×10^−9^; FM, FMT, p=0.0036; W, p=0.40; WS, p=0.11; Rw, p=0.22; T, p=0.0012. Right: multiple comparison between feature groups were plotted, pooling data from both layers.

We fit our model with LASSO to automatically identify relevant factors influencing each neuron’s responses. We clustered neurons into different feature sensitive groups by identifying the predictors with the maximum ΔR^2^ within each neuron, calculated by shuffling predictor values (Figure 2B, see Method). We visualized the distribution of these groups of neurons in a lower dimension space projected by T-sne (Supplemental Figure 1A), which verified that such identified groups of neurons segregate in the projected space. These results suggest that A1 neurons encode various features in addition to sound during the behavioral session. For example, Rw neurons were robustly and preferentially activated during reward consumption (Supplemental Figure 1B) while showing little sensitivity to other features (Figure 2B, Rw).

### Action driven activity is represented in more neurons in L2/3 than in L4

A1 is thought to perform hierarchical computations (Winkowski and Kanold, 2013) and thus the fraction of neurons encoding non-auditory variables might differ between layers. Thus, we quantified the fraction of neurons within each FOV that were assigned to each group as a function of layer (Figure 2C). Among all groups, F and FT neurons were the most abundant, while FM and FMT neurons were sparser, suggesting that A1 neurons are more sensitive to the presence of sound energy within its receptive field, consistent A1 neurons being more responsive to spectrally static stimuli than to frequency modulations (Issa et al., 2017). Across cortical layers, L4 had more neurons assigned to F and FT group than L2/3 while the two layers showed similar proportions of FM and FMT neurons. In contrast, L2/3 showed more abundant W and WS neurons, suggesting a higher degree of integration of action related information in L2/3.

A1 neurons show diverse tuning properties (Bandyopadhyay *et al*., 2010; Liu and Kanold, 2021; Maor et al., 2016; Rothschild *et al*., 2010; Sutter et al., 1999) and it is possible that action integration depended on sound tuning properties. We thus investigated the tuning properties of these feature sensitive groups of neurons (Figure 2D, E). Sound encoding neuronal groups, i.e., F, FT and FM, FMT, tend to show more neurons responsive to tones (Figure 2D) and showed broader tuning than other neuronal groups (Figure 2E). In addition, sound encoding L2/3 neurons showed broader tuning than L4 neurons, consistent with previous reports (Bowen et al., 2020). Together, these results suggest that there are functionally separate groups in A1 that encode sound or action related information and that there is enhanced encoding behaviorally relevant information in L2/3.

### Sound driven responses in A1 were more suppressed during behavior but suppression varied by layer

A1 is in a more suppressed state during behavior, i.e., A1 neurons’ responses to tones are weaker during locomotion, licking or task engagement than during passive listening (Bigelow et al., 2019; Kuchibhotla et al., 2017; Otazu et al., 2009; Rummell *et al*., 2016; Zhou et al., 2014). Although not a key focus of the current study, we also confirmed these observations by investigating the coefficients of sound information encoding variables (F and FM) and their corresponding interaction term with the task variable (FT and FMT, Supplemental Figure 2) in frequency (F and FT) and frequency modulation (FM and FMT) sensitive neurons. The signs of the respective coefficient pair, e.g., F and FT term, determine the relative strength of neuronal responses to sound across active and playback sessions (Supplemental Figure 2A, B). The majority of F, FT and FM, FMT neurons showed weaker responses to the preferred sound features in the active session than in the playback session (Supplemental Figure 2C, D). Although other forms of active-playback interaction exist, e.g., stronger responses in the active session than in the playback session (Supplemental Figure 2A, B), the suppressive effect of active session to sound responses were the strongest among all forms of active-playback interactions (Supplemental Figure 2E, F). In addition, the suppressive effect was stronger in L4 than in L2/3 for F and FT neurons (Supplemental Figure 2B, C) while the same phenomenon was absent for FM and FMT neurons (Supplemental Figure 2E, F). Together, our model reaffirmed previous findings of behavior modulation of sound responses in A1, further validating our modeling approach.

### A1 neurons showed wheel-turning driven activity that was turn direction selective and sound dependent

Previous studies have consistently shown that in A1, the actions of the animal influence sound encoding, and thus action related input is integrated in the A1 (Clayton et al., 2021; Rummell *et al*., 2016; Schneider *et al*., 2018). Similarly, our model identified a group of A1 neurons sensitive to the wheel turning action of the mice (Figure 2B, W and WS). We first examined how sounds and action were integrated in W neurons. Figure 3A shows an example W neuron from the same group shown in Figure 2B. The activity of this neuron was temporally correlated with leftward wheel turning and this correlation existed both within trials when the task sound was presented and during intertrial intervals when no task sound was presented. Furthermore, this neuron’s responses tended to be smaller during trials than during intertrial intervals (Figure 3A, arrows and arrow heads, respectively). This example neuron’s activity was preferentially corelated with leftward wheel turning but not with rightward turning, suggesting turn direction selectivity in W neurons. Thus, we separated W neurons into right turn and left turn preferring neurons by identifying which specific turn direction term (+W for right turns and - W for left turns) had a higher ΔR^2^ and thus better explained the variability in the neuron’s responses. Right turn preferring neurons had prominently positive +W coefficients while left turn preferring neurons had prominently positive -W coefficients (Figure 3B, C). Furthermore, the sign (positive or negative) of the interaction term between wheel turning and task sound presentation, i.e., WS, would inform whether the task sound facilitated or suppressed wheel turning induced neuronal responses. We found that right turn coefficient pairs (+W and +WS) and left turn coefficients pairs (-W and -WS) were negatively correlated in right and left turn preferring neurons respectively, suggesting that the W neurons showed a suppressed action driven responses during sound presentations (Figure 3B, C). Moreover, the absolute values of WS coefficients were smaller than the values of W coefficients, indicating that the WS term is modulatory. These action related information within A1 could be interpreted as the input of the forward model in A1, which potentially encodes the predicted sound from the underlying action.

**Figure 3.**
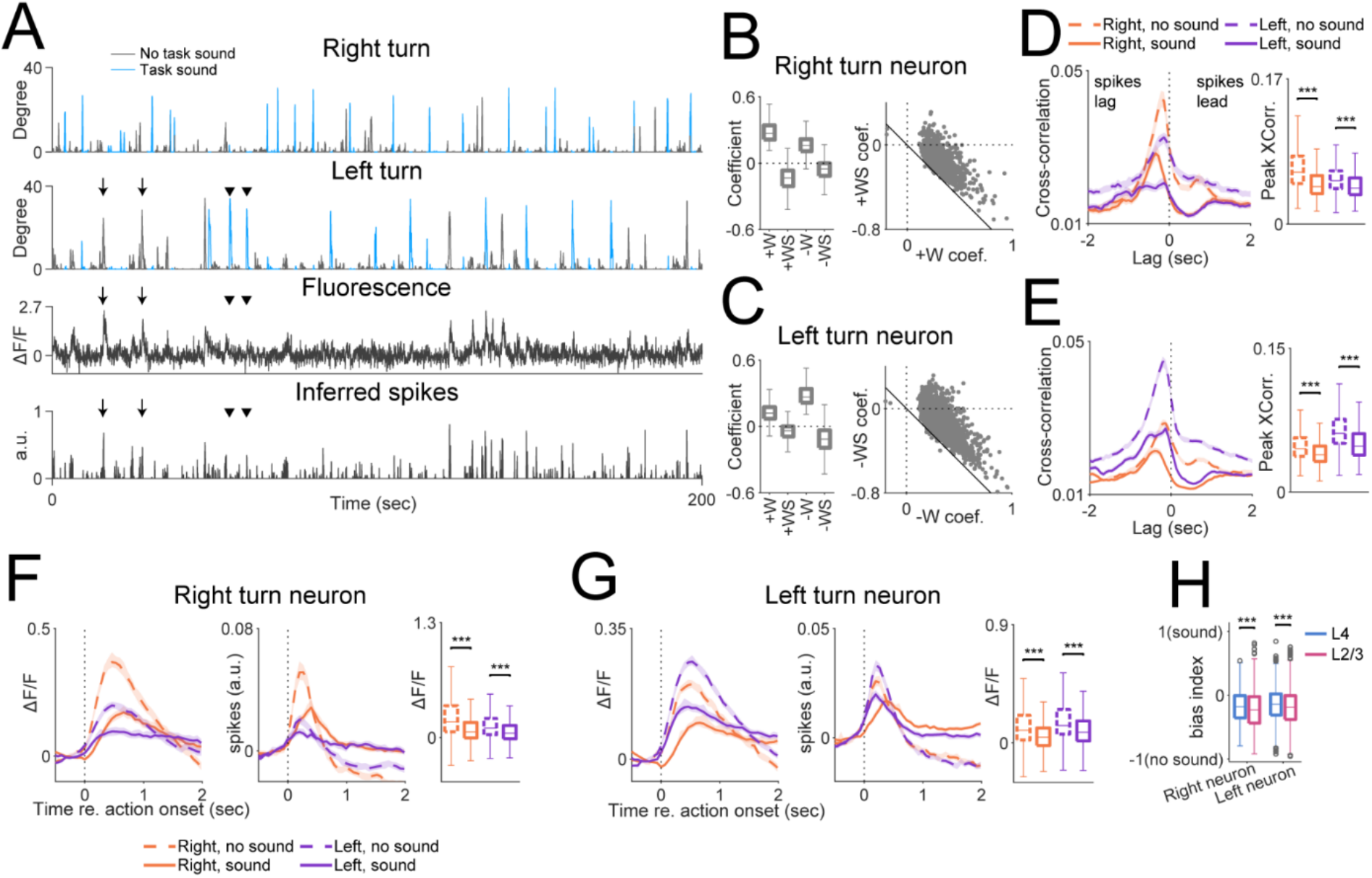
Action sensitive (W) neurons show sound dependent activity. (A) Top two rows: example left- and right-turning wheel movements are plotted. Wheel movements during sound presentations are plotted in orange. Inter-trial wheel movements are plotted in blue. Bottom two rows: fluorescence trace and inferred spikes from one example action sensitive (W) neuron from the same behavioral session. This neuron was preferentially sensitive to left turns, and its response amplitude tended to be smaller during sound presentations (arrow heads) than during inter-trial intervals (no sound, arrows). (B) Left: boxplot showing the coefficients of right turn preferring W neurons. In these neurons, +W and +WS term coefficients are more prominent than the negative counterparts. Right: scatter plot showing the joint distribution of +W and +WS coefficients. +W and +WS coefficients had a significantly negative correlation (p=1.5×10^−228^). The black solid line is of slope −1. (C) The same as in (B) but for left turn preferring W neurons. For these neurons, -W and -WS coefficients also had a significantly negative correlation (p<10^−4^). (D) Left: the correlogram between inferred spikes of right turn preferring W neurons and the onset of wheel movements broken down by both turn directions (right vs. left) and sound presence (sound vs. no sound). Right: boxplot showing the peak value of the correlogram under the four conditions. The peak correlogram values were higher during no sound period than during sound presentation for both right and left turns (Wilcoxon sign rank test, right, no sound vs. right, sound, p=2.1×10^−74^; left, no sound vs. left sound, p=2.9×10^−32^). (E) Same as in (D) but for left turn preferring W neurons. Similarly, the peak correlogram values were higher during no sound period than during sound presentation for both right and left turns (Wilcoxon sign rank test, right, no sound vs. right, sound, p=1.6×10^−91^; left, no sound vs. left sound, p=1.7×10^−152^). (F) Left and middle: the fluorescence and inferred spike traces of right turn preferring W neurons, temporally aligned to the onset of wheel turning and broken down by the turning directions and the presence of sound. Right: boxplot showing the ΔF/F following the wheel turning onset, broken down by the four conditions. ΔF/F values following wheel turning were higher during no sound than during sound presentation (Wilcoxon sign rank test, right, no sound vs. right, sound, p=1.0×10^−68^; left, no sound vs. left sound, p=2.1×10^−33^). (G) Same in (F) but for left turn preferring W neurons. Similarly, ΔF/F values following wheel turning were higher during no sound than during sound presentation (Wilcoxon sign rank test, right, no sound vs. right, sound, p=5.9×10^−107^; left, no sound vs. left sound, p=8.5×10^−96^). (H) Boxplot showing the bias indices quantifying the strength of the suppressive effect of sound to neuronal responses evoked by wheel turning, broken down by neuron groups (left or right turn preference) and cortical layer. More negative values suggest stronger suppressive effects. For both right and left turn preferring neurons, the suppression strength was higher in L2/3 than L4 (Wilcoxon sign rank test, right preferring neuron, p=4.3×10^−4^; left preferring neuron, p=6.2×10^−8^).

As our model had a coarse temporal resolution (0.5 second time window), we further verified the suppressive effect of the task sound on W neurons by two separate analyses with the same temporal resolution as the imaging frame rate. First, we constructed correlograms between W neurons’ inferred spikes and the onset of wheel movement (Figure 3D, E). For both left and right turn preferring neurons, the correlations were largest between spikes and wheel movement during the period with no task sound, suggesting that neuronal activities were more time-locked to wheel movement in quietness (Figure 3D, E). Furthermore, the negative lags of the peaks of the correlogram indicate that the onset of wheel movement preceded inferred spikes, suggesting that neuronal activities were driven by the wheel movement (Figure 3D, E). Second, we examined the neuronal responses temporally aligned to the onset of wheel movement, broken down by turning directions and the presence of sound stimuli (Figure 3F, G). This analysis shows that turning in the preferred direction during no-sound intertrial period evoked the largest responses in both left and right turn preferring neurons. Together, these analyses show that wheel movements in quietness evoked responses with a higher temporal coherence and a larger amplitude in W neurons, confirming the suppressive effect of task sound on W neurons. Furthermore, the suppressive effect of task sound is also consistent with the interpretation that W neuron’s responses potentially represent predicted sound from action. In the absence of the actual task sound, the responses were larger due to the deviance from the expected sensory outcome, while the responses were smaller when matching task sound was presented.

As L2/3 showed more action selective neurons than L4 (Figure 2C), we speculate that the degree of suppressive effect of sound in W neurons could also show hierarchical layer dependence. Thus, we computed a suppression index using the response amplitude following wheel movement onset during sound presentation (R_sound_) or in quietness (R_no-sound_), i.e., (R_sound_-R_no-sound_)/(R_sound_+R_no-sound_) and compared the index values across L2/3 and L4. This analysis revealed that L2/3 W neurons showed a stronger sound induced suppressive effect than L4 W neurons (Figure 3H), which suggests that while L4 receives feedforward sound information, L2/3 represents a locus for the interaction between output of the forward model (predicted sound) and the actual sound. Together, our analysis further shows that the responses of these action neurons are dynamically modulated by the presence of sound, which could provide a substrate for more complex associations between sound and action within A1.

### A subset of A1 neurons is jointly sensitive to sound and action and encode the spectral contingencies of the task

In addition to the action neurons (W neurons) identified above, we also identified a group of neurons selective to the WS feature, i.e., the interaction between the wheel movement and the sound presentation, suggesting that these neurons are jointly sensitive to both action and sound (Figure 2B). A closer inspection of the covariation between action and sound during a trial reveals that a typical trial (e.g. Figure 2A) consisted of not only periods where the carrier frequency changes following the wheel movement, but also periods the carrier frequency decoupled from the wheel movement at the spectrum boundary (Figure 1E, 2A, 4A). Due to the design of our interactive behavior paradigm, the carrier frequency cannot increase beyond the upper boundary of the spectrum (40 kHz) or decrease below the lower boundary of the spectrum (10 kHz). Thus, any wheel turning beyond these spectrum boundaries would essentially produce no carrier frequency change and will result in a decoupling between the action and sound (DCP, Figure 2A, 4A). Thus, there is a mismatch between the predicted change in sound frequency and the actual clipping of sound frequency at the boundaries. These DCP events happen within WS periods, i.e., wheel turning during sound presentations, and thus DCP was closely temporally related to WS (Figure 4B). The correlogram shown in Figure 4B suggests that the DCP at the spectral boundary typically followed the wheel movement onset with a lag of 0.2 second, which was within the time window chosen for our linear model (0.5 second). Therefore, it is possible that a subset of WS neurons identified in Figure 2B was sensitive to these DCP events.

**Figure 4.**
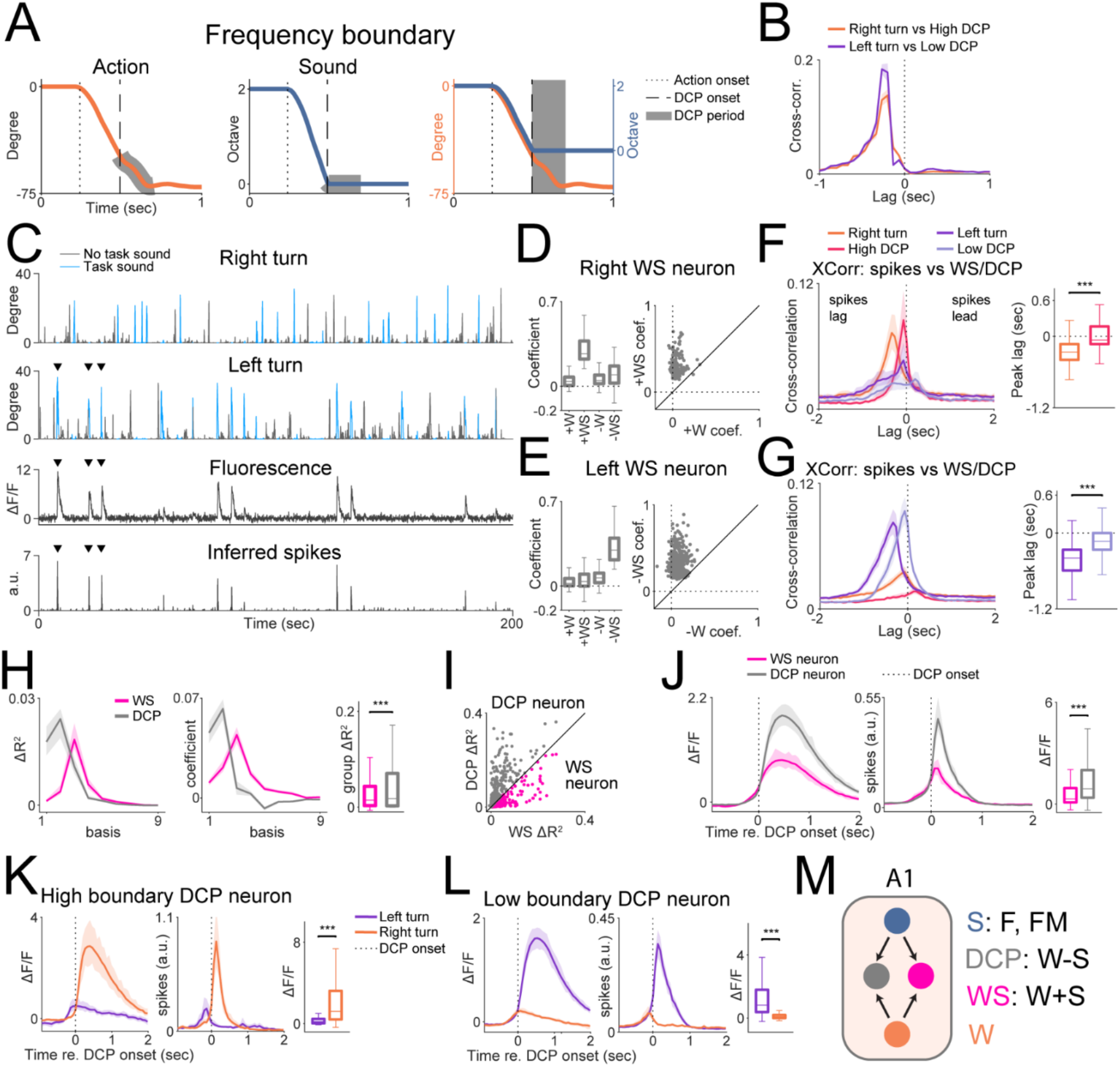
A subset of action-sound sensitive neurons encode the spectrum boundary of the task paradigm. (A) The wheel (left) and frequency (middle) trajectory of an example trial. The decoupling (DCP) period where the wheel movements were not translated into carrier frequency changes due to the spectrum boundary was labeled in gray in both plots. The right plot highlights the disassociation of the action and the sound at the spectrum edge. The dotted line marks the onset of the action while the dashed line marks the onset of the DCP period. (B) The correlogram between onset of left turn events and the onset of DCP at the low frequency boundary (Low DCP, purple curve) and the correlogram between onset of right turn events and the onset of DCP at the high frequency boundary (High DCP, orange curve) are shown. Both correlograms peaked at around −0.2 second. The solid lines represent the mean, while the shaded regions represent standard deviation. (C) Top two rows: example left- and right-turning wheel movement from one active session. Wheel movements during sound presentations are plotted in orange. Inter-trial wheel movements are plotted in blue. Bottom two rows: fluorescence trace and inferred spikes from one example action-sound jointly sensitive (WS) neuron from the same behavioral session. This neuron was preferentially activated by left turns during trials (arrow heads). (D) Left: boxplot showing the coefficients of right turn preferring WS neurons. In these neurons, the +WS term had the most prominent coefficients. Right: scatter plot showing the joint distribution of +W and +WS coefficients. +WS coefficients were higher than +W coefficients (Wilcoxon rank sum test, p=9.6×10^−25^). The black solid line has a slope of 1. (E) The same as in (D) but for left turn preferring WS neurons. Similarly -WS term coefficients were larger than -W coefficients (Wilcoxon rank sum test, p=1.7×10^−69^). (F) Left: the correlogram between inferred spikes of right turn preferring WS neurons and the onset of either wheel movement (left or right turn) or DCP events (Low DCP or High DCP). Spiking activities had a shorter lag from High DCP events than from right turn onset (Wilcoxon sign rank test, p=9.2×10^−11^). (G) Same as in (F) but for left turn preferring WS neurons. Spiking activities had a shorter lag from Low DCP events than from left turn onset (Wilcoxon sign rank test, p=2.6×10^−12^). (H) Left and middle: the ΔR^2^ and coefficients as a function of the 9 bases plotted separately for WS and DCP predictors. For WS predictors, the 3^rd^ basis had the most explanatory power, while for DCP predictors, the 1^st^ and 2^nd^ basis had the most explanatory power. Right: boxplot showing the group ΔR^2^ from all bases plotted as a function of WS and DCP. DCP predictors had higher ΔR^2^ than WS predictors (Wilcoxon sign rank test, p=2.2×10^−6^). (I) The scatter plot showing the joint distribution of group ΔR^2^ of WS and DCP term. We identified the putative DCP sensitive neurons as the subset of neurons with higher DCP group ΔR^2^. The solid line has a slope of 1. (J) Left and middle: the fluorescence and inferred spikes of WS and DCP neurons as grouped in (H) were temporally aligned to the onset of DCP and plotted. DCP neurons showed larger responses following DCP onset compared to WS neurons (right, Wilcoxon rank sum test, p=1.3×10^−15^). (K) Left and middle: fluorescence and inferred spike traces of high frequency boundary DCP preferring neurons, temporally aligned to the DCP onset. These neurons had larger ΔF/F responses to right turn induced DCP at the high frequency boundary (right plot, Wilcoxon sign rank test, p=1.0×10^−9^). Vertical dotted lines indicate the onset of DCP events. (L) Left and middle: same as (K) but for low frequency boundary DCP preferring neurons. These neurons had larger ΔF/F responses to left turn induced DCP at the low frequency boundary (right plot, Wilcoxon sign rank test, p=2.0×10^−39^). (M) While DCP-insensitive WS neurons could represent the facilitative responses due to matching sound and the predicted sound constructed by the forward model using information represented by W neurons (W+S), DCP neurons encode the deviance between the two (W-S).

Figure 4C shows an example WS neuron, and it is evident that this neuron was selective to left turning wheel movements during sound presentations. We thus further divided WS neurons into right turn and left turn preferring subgroups (Figure 4D, E), with further analyses performed separately on these two subgroups. First, to explore the possibility that WS neurons were sensitive to DCP events at the spectral boundaries, we examined the correlogram between WS neurons’ inferred spikes and wheel movement onset and the correlogram between spikes and the onset of DCP events (Figure 4F, G). For both left and right preferring WS neurons, their inferred spikes had shorter lags from DCP onset than from movement onset, which suggests that these responses could be more directly driven by DCP events. Next, we used a model with higher temporal resolution to address this question (Supplemental Figure 3, see also Method). We fit the WS neurons’ inferred spikes to these predictors and examined their explanatory powers (Figure 4H). For WS predictors, the 3^rd^ basis had the highest ΔR^2^ while for DCP predictors, the 1^st^ and 2^nd^ basis had the highest ΔR^2^. As these bases have peak activations that are progressively delayed in time (the 1^st^ basis has the earliest activation), the fact that WS neurons’ responses were better explained by the first two bases convolved with the DCP events suggest that WS neurons’ responses were more temporally aligned with DCP onset. In addition, DCP predictors had a larger overall ΔR^2^ than WS predictors (Figure 4H), suggesting that DCP could better explain the responses. We assigned the subset of neurons that had higher DCP ΔR^2^ than WS ΔR^2^ as DCP neurons while the rest as WS neurons (Figure 4I). Next, we examined the fluorescence and inferred spike traces of both DCP and WS neurons, temporally aligned to the DCP onset (Figure 4J). DCP neurons had a larger increase in ΔF/F following the DCP onset than WS neurons (Figure 4J). Furthermore, DCP neurons had a sharp increase in firing rate following DCP onset, while WS neurons’ firing rate largely decreased following DCP onset (Figure 4J, middle). Together, these results indicated that WS neurons constituted three subgroups: neurons insensitive to DCP events and two groups of neurons sensitive to low and high boundary DCP events respectively. The responses of the identified DCP neurons could reflect error signals, as the relationship between action and carrier frequency changes breaks down at the spectrum boundary. Furthermore, DCP neurons responded selectively to low and high frequency boundary DCP events (Figure 4K, L). These results together suggest that a subset of sound and action jointly sensitive neurons responded robustly to the DCP of action and frequency at the spectrum boundary. We speculate that these DCP neurons help guide the behavior of the animal to stop wheel turning as these DCP neurons signal reaching the target frequency zone. Furthermore, as suggested in the previous section that W neurons’ responses could be interpreted as part of the forward model representing the predicted sound, the WS neurons’ responses could also be interpreted as the interaction between the predicted and the actual sound. While DCP-insensitive WS neurons represent a facilitated responses when the predicted and actual sound match, the DCP sensitive neurons encode the deviance between the two (Figure 4M).

### Delay Sound (DS) perturbations induced both action driven and sound driven error responses in subsets of A1 neurons

The identification of DCP sensitive neurons indicate that a subset of A1 neurons can signal errors induced by the mismatch between action and carrier frequency change. However, the DCP at the boundary of the spectrum is intrinsic to the task design and thus we hypothesize that such DCP could be expected or learned by the animal. We thus sought to investigate if A1 also encode unexpected violations of the action-sound relationship which could generate error signals. For this purpose, we introduced two distinct perturbations: “Delay Sound” (DS) and “Stop Frequency” (SF) (Figure 1E). DS perturbations aimed to perturbate the timing relationship between action and sound (perturbate “when”) while SF perturbations aimed to perturbate the sensory feedback (perturbate “what”).

Figure 5A shows an example DS trial. We introduced a delay (Δt) of 1 second between the action and the carrier frequency change. Due to the delay, the carrier frequency remained same throughout the first 1-second period. Thus, any action during this period triggered an action induced DCP event (DCP_A_). After the 1-second delay, carrier frequency started changing, but as it was not matched to the concurrent action, this triggers another DCP event but induced by sound (DCP_S_). DCP_A_ and DCP_S_ were thus temporally separate DCP events driven by different features. In order to identify error responses triggered by DCP_A_ or DCP_S_, we built a second linear model focused on the 2 seconds after the action onset (Figure 5A). We constructed the predictors from 4 non-overlapping time windows of 0.5-second duration (Figure 5B). The first two windows fell within the first 1 second, which we denoted as the action window, while the other two windows fell within the 2^nd^ 1 second, which we denoted as the sound window. Each trial thus contributed 4 data points to the model. In addition to DS trials, we also included non-DS (NDS) trials and playback of DS trials (DSPB). We selected DS and NDS trials that resulted in hits where the animals made significant wheel movement, which was comparable across DS and NDS trials and resulted in similar frequency trajectories despite the delay of sound in DS trials (Figure 5C). The playback of DS trials or DSPB trials shared the same frequency trajectories as DS trials, but the animal did not turn the wheel (Figure 5C, see also Figure 1D). These trials thus allowed us to distinguish the contribution of factors other than DCP_A_ or DCP_S_ to the neuronal responses.

**Figure 5.**
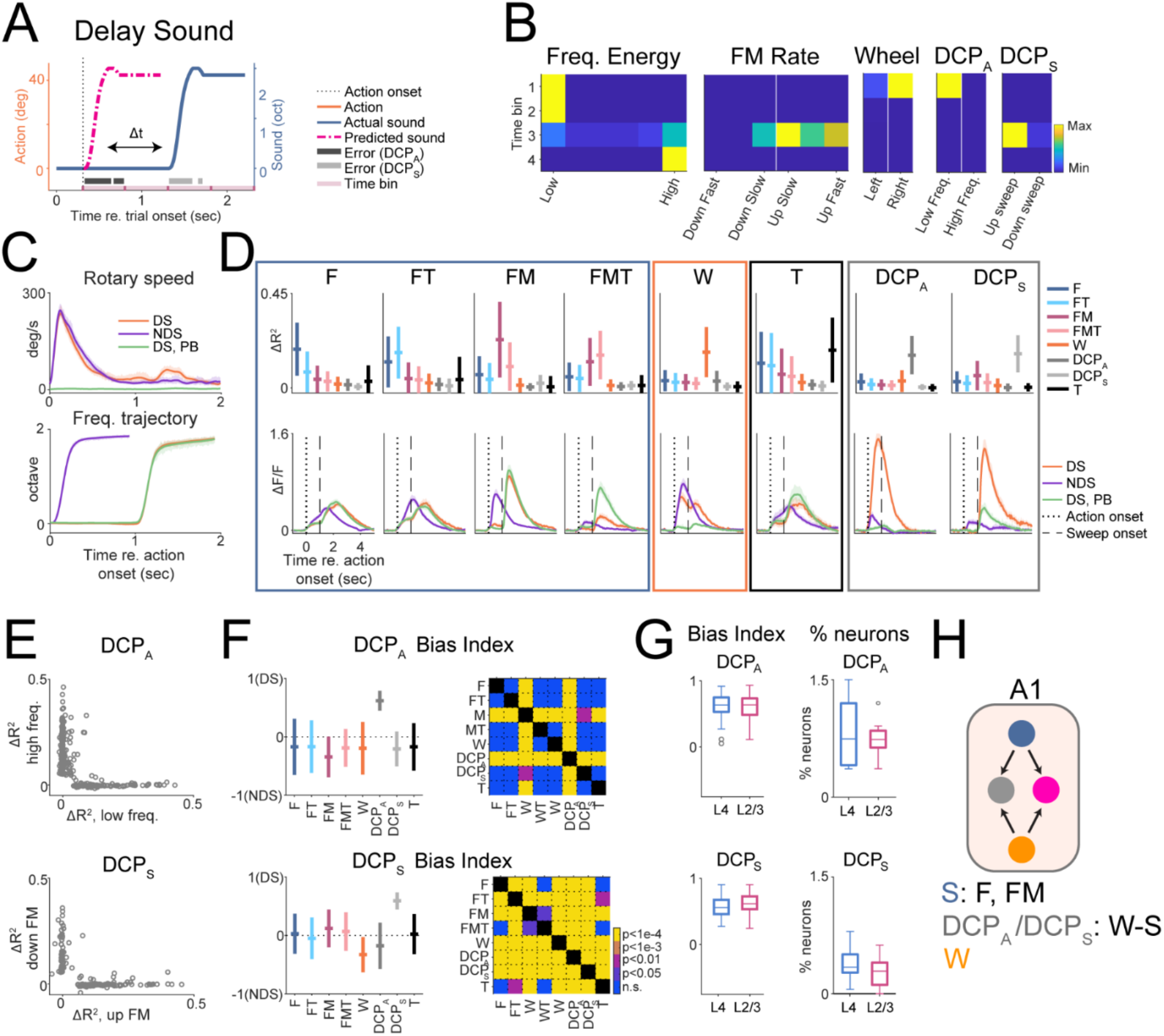
“Delay-Sound” (DS) perturbation induced error responses in a subset of A1 neurons. (A) The frequency (blue solid curve) and wheel trajectory (orange solid curve) of one example DS trial are shown. The violet-red dotted-dash curve shows the “would-be” or predicted frequency trajectory without the DS perturbation. Two forms of DCP (action induced DCP or DCP_A_ and sound induced DCP or DCP_S_) are marked in dark and light gray, respectively. Four windows of 0.5 second duration (marked on top of the plot in purple) were placed following the onset of the action to extract the predictors as shown in (B). (B) The predictors extracted from the example trial in (A). (C) Top: average absolute wheel speed as a function of trial conditions. The wheel speed of DS and NDS trials were similar from 0 to 1 second, resulting in similar frequency trajectories despite the 1-second delay as shown in the right plot. DS: delay-sound trials; NDS: non-delay-sound trials; DSPB, playback of delay-sound trials. Bottom: the corresponding absolute frequency trajectories of the three trial conditions. (D) Top: feature sensitive neurons were grouped according to their maximum ΔR^2^ term. The horizontal lines represent the mean while the vertical lines represent standard deviation. Bottom: the average fluorescence traces from the corresponding feature sensitive neuronal groups as a function of trial conditions. The solid curves represent the mean while the shaded regions represent 95% confidence intervals. The dotted vertical lines mark the action onset while the dashed line mark the onset of the delayed frequency sweep. (E) Scatter plots showing the joint distribution of ΔR^2^ of the low frequency and high frequency term for DCP_A_ neurons (top) and the joint distribution of ΔR^2^ of the up FM sweep and down FM sweep term for DCP_S_ neurons. Both groups of DCP neurons showed a high selectivity for a single feature. (F) Top left: DCP_A_ bias index as a function of different feature sensitive groups shown in (D). The more positive the value of the bias index is, the higher the strength of the action induced error signal. Top right: the multiple comparison between the different feature sensitive groups. Bottom left: DCP_S_ bias index as a function of different feature sensitive groups shown in (D). The more positive the value of the bias index is, the higher the strength of the sound induced error signal. Bottom right: the multiple comparison between the different feature sensitive groups. (G) Left: boxplot showing the DCP_A_ and DCP_S_ bias indices as a function of cortical layer within DCP_A_ and DCP_S_ neurons, respectively. DCP_A_ bias index showed no difference across layers (Wilcoxon rank sum test, p=0.42). DCP_S_ bias index also showed no difference across layers (Wilcoxon rank sum test, significance adjusted with Bonferroni correction, p=0.032). Right: boxplot showing the percentage of neurons in a FOV that were assigned to either DCP_A_ or DCP_S_ groups as a function of cortical layer. There was no difference across layers (L4 vs. L2/3, Wilcoxon rank sum test, DCP_A_, p=0.89; DCP_S_, p=0.26). (H) Similar to Figure 4M, DCP_A_ and DCP_S_ neurons likely represent the deviation between the predicted sound and the actual sound feedback.

We fit a model to the neuronal responses with features including F, FM, W as well as DCP_A_ and DCP_S_, and we identified different feature sensitive neurons using a similar approach as in Figure 2B. The average fluorescence traces from each group of neurons showed distinct patterns and we identified 2 new groups of neurons as their responses could be explained by DCP_A_ or DCP_S_ (Figure 5D). Other feature sensitive neurons did not show selectivity toward DCP_A_ or DCP_S_ events (Figure 5D). Rather, their responses were time-locked to their preferred features. For example, wheel movement preferring neurons (W neurons) responded similarly following wheel movement onset regardless of whether the sound was delayed (Figure 5D, W neurons, DS vs. NDS). In contrast, DCP_A_ neurons showed larger responses in the action window during DS trials than during NDS trials, while DCP_S_ neurons showed larger responses in the sound window during DS trials than during NDS trials or during DPSB. As these responses could not be fully explained by only considering the frequency content of the sound (DS trials vs DSPB trials) or the animal’s action (DS trials vs NDS trials), they likely encode error signals induced by the mismatch between the action and the sound feedback.

As DS was introduced in both trials with low and high starting frequencies, we investigated whether the responses of DCP_A_ and DCP_S_ were also selective to the acoustic context (Figure 5E). For DCP_A_ neurons, their responses were explained mostly by either DCP at the low starting frequency (resulted from right turns) or at the high starting frequency (resulted from left turns). For DCP_S_ neurons, their responses were evoked by either up frequency sweep or down frequency sweep. Thus, these error responses not only encoded the presence of DCP events, but they also encoded the specific sound content of the DCP events.

One of the key questions of the current study is to investigate whether the strength of the error signals would have hierarchical structures across processing stages such as cortical layers. However, comparing the amplitude of ΔF/F could be biased due to the difference of signal to noise ratio at different imaging depth. We thus quantified the strength of these error signals by computing a bias index that alleviates such biases through internal normalizations. Specifically, for DCP_A_ induced errors, we computed for each neuron the bias index between responses from the action window in DS trials (R_DS_) and responses from the action windows in NDS trials (R_NDS_), i.e., (R_DS_-R_NDS_)/(R_DS_-R_NDS_). Similarly, for DCP_S_ induced errors, we computed the bias index using the responses from the sound window in DS trials and responses from the action window in NDS trials. Among all feature sensitive neurons, DCP_A_ and DCP_S_ neurons showed the most positive bias indices and thus showed the strongest error responses as expected from their temporal responses (Figure 5F). We proceeded to compare the same bias indices across L4 and L2/3 in A1 but we did not find a difference in either DCP_A_ or DCP_S_ error signal strength across the two cortical layers (Figure 5G, left). Moreover, the fraction of neurons in L4 and L2/3 that showed either DCP_A_ or DCP_S_ induced responses was also similar (Figure 5G, right). Overall, these results suggest that specific A1 neurons represented distinct error signals and that error signals were similarly represented in L4 and L2/3 of mouse A1. Similar to the DCP neurons sensitive to the frequency boundary (Figure 4M), DCP_A_ or DCP_S_ neurons’ responses could also be interpreted as encoding the deviation of the actual sound from the predicted sound, whose construction relied on the output of the forward model that utilized the action related information represented by the W neurons (Figure 5H).

### “Stop-Frequency” (SF) perturbations evoked frequency specific error responses in a subset of A1 neurons

The DS perturbations disturbed the temporal relationship between the sound and the action (perturbate “When”). To directly perturbate the content of the sound feedback (perturbate “what”), we introduced a second form of perturbation, i.e., “Stop-Frequency” (SF) perturbations. In SF trials, as the carrier frequency approached one of the four SFs, randomly chosen for a given trial, the frequency stream would be ‘frozen’ for a brief period (0.25 second) at the SF and thus introducing a frequency dependent DCP (Figure 6A). This brief action driven DCP created a deviation from the predicted frequency trajectory, and we hypothesized that A1 neurons also encode DCP at these SFs as error signals.

**Figure 6.**
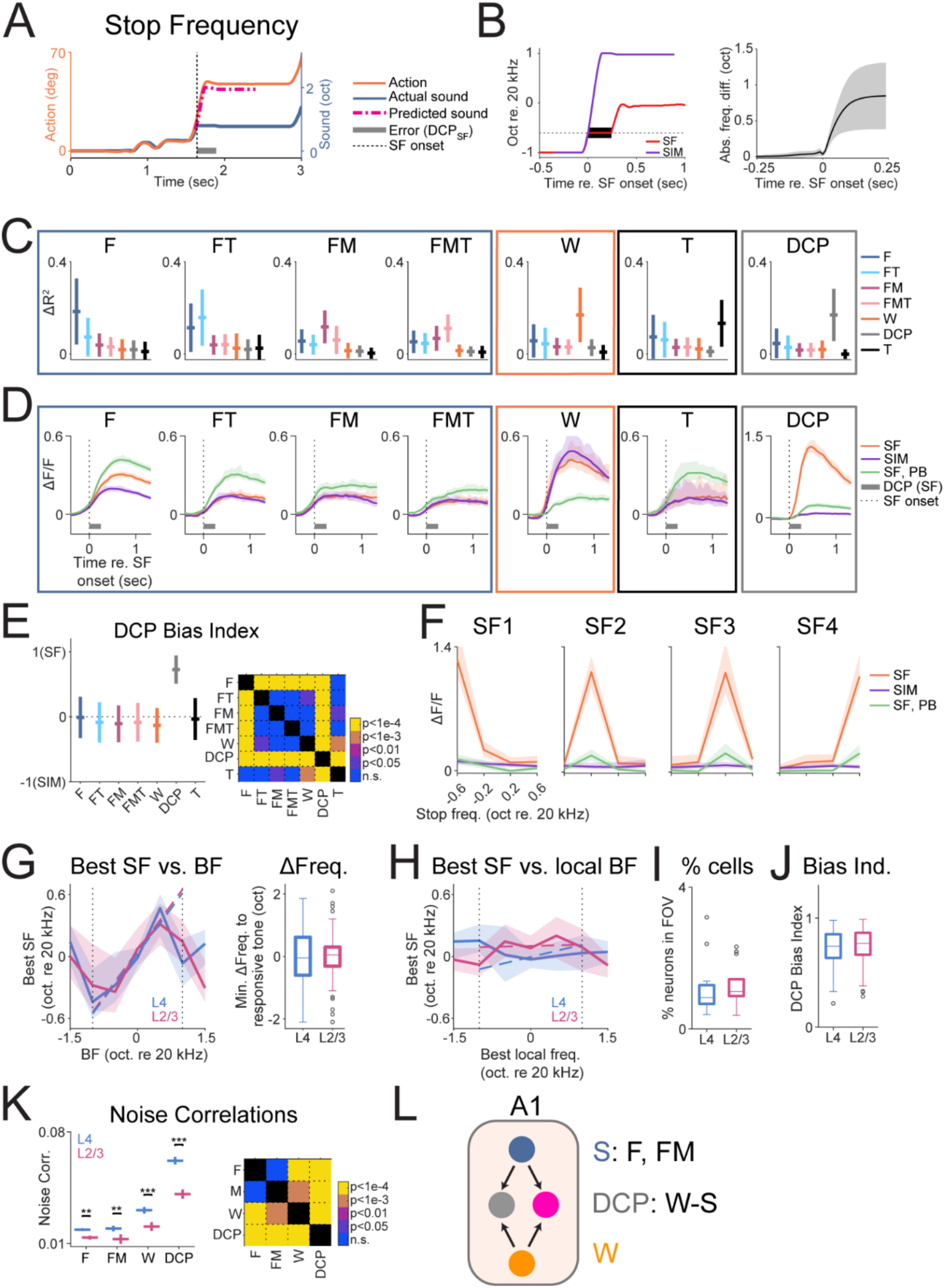
“Stop-frequency” (SF) perturbations induced DCP responses at frequencies outside target zones. (A) Example frequency and action trajectories of an example SF perturbation trial. The orange solid line shows the frequency trajectory, and the gray horizontal line marks the period of the SF perturbation, where the carrier frequency was frozen despite the wheel movement. The violet-red dash-dotted line shows the predicted frequency trajectory if the perturbation had not been introduced. The blue line shows the corresponding wheel movement of the trial. (B) Left: the frequency trajectory of an example active trial with perturbation (red) and the frequency trajectory of an example SIM trial (purple) that shared a similar frequency trajectory to the SF trial immediately before DCP onset (marked in black). Right: the average absolute difference in frequency trajectories between SF trials and SIM trials. The difference was negligible before the perturbation onset and increased after perturbation onset. (C) The group ΔR^2^ of identified feature sensitive neurons are shown. (D) Average ΔF/F traces from SF, SIM and playback (PB) trials from the corresponding feature sensitive neurons as shown in (C). Solid lines show average across neurons while the shaded regions show 95% confidence interval. The vertical dotted lines indicate the onset of SF perturbation. The black bar indicates the duration of the SF perturbation. (E) Left: the bias indices quantifying the difference between SF and SIM trials were plotted as a function of feature sensitivity. Right: the multi-comparison table between the groups is shown. (F) DCP_A_ neurons showed distinct preference for SFs and neurons could be categorized by the SF that evoked the largest response. Each panel shows the average ΔF/F responses over the subpopulation of neurons with the same SF preference. (G) Left: for neurons showed passive tone responses, the SF that each DCP neuron was most sensitive to (Best Stop Frequency, BSF) was plotted against the neuron’s best frequency (BF). BSF and BF were positively correlated within the carrier frequency boundary (10 and 40 kHz, marked by the dotted lines). The linear fit was plotted in dash lines (L4, slope=0.58, p=2.9×10^−5^; L2/3, slope=0.59, p=3.0×10^−7^). Right: for the same subset of tone responsive DCP neurons, the distance of BSF to the nearest frequency that evoked significant responses was shown as a function of cortical layer. Both distributions centered around zero. (H) Local BF (LBF), the frequency evoking the most responses from neurons within a radius of 100 μm of DCP neurons, was plotted against the DCP neurons’ BF. LBF was not correlated with BSF in DCP neurons (L4, slope=0.12, p=0.11; L2/3, slope=0.01, p=0.87). (I) Boxplot showing the bias indices from DCP_A_ neurons as a function of cortical layer. No difference was found between L4 and L2/3 (Wilcoxon rank sum test, p=0.080). (J) Boxplot showing the fraction of neurons per FOV identified as SF responsive as a function of cortical layer. No difference was found between L4 and L2/3 (Wilcoxon rank sum test, p=0.26). (K) Left: cross-group noise correlation (NC) between DCP and F, FM, W neurons (column 1-3) and intra-group noise correlation within DCP neurons (4^th^ column). Vertical lines show the mean and the vertical lines show the SEM. L4 vs. L2/3, Wilcoxon rank sum test, NC DCP and F p=0.0012; NC DP and FM, p=0.0058; NC DCP and W, p=2.6×10^−5^; NC DCP vs DCP, p=4.6×10^−11^. Right: the multiple comparison table showing the difference of NC regardless of cortical layer. (L) The responses of SF induced DCP neurons can be again interpreted as the deviation between the predicted sound generated by the forward model in A1 using the action related information from the actual sensory input (W-S).

To identify the neurons that were sensitive to DCP at these four SFs, we constructed a similar model to that of the DS perturbation but focused on the time immediately before and after the onset of the SF (see Method). Both SF trials and the corresponding playback trials were included, such that the contribution of sound feature selectivity to the responses could be accounted for. We also included non-perturbated active trials where the frequency trajectories were similar to the perturbated trials immediately before the SF onset (Figure 6B, SIM trials). The similarity of the frequency trajectory and the presence of wheel movement allowed us to account for the contributions of both action and sound feature selectivity to the neuronal responses. We hypothesize that true DCP responses are specific to SF trials and thus DCP sensitive neurons should show weaker responses in SIM and playback trials than in SF trials.

We thus fitted a linear model using LASSO to the neuronal responses and were able to identify neuronal groups that were sensitive to different features (Figure 6C). Next, we examined the response profile of these groups of neurons (Figure 6D). Except for DCP neurons, all other groups of neurons showed rather similar responses in SF and SIM trials, suggesting a lack of selectivity to DCP. In F, FT, FM and FMT neurons, playback trials tend to evoke larger responses, confirming a more suppressed state in active trials (see Figure 2). W neurons showed similar responses in SF and SIM trials, with little activation in playback trials, confirming their sensitivity to actions. Moreover, these groups of neurons showed responses before DCP onset and thus were driven by factors other than DCP. In contrast, DCP neurons showed responses after DCP onset while showing much weaker responses in both SIM and playback trials. Thus, DCP neuron’s responses cannot be accounted for by stimulus selectivity as well as the animal’s action, and thus likely represent bona fide error responses. We quantified the strength of the error responses by computing a bias index between the response amplitude of the SF trials (R_SF_) and the response amplitude of the SIM trials (R_SIM_), i.e., (R_SF_ - R_SIM_)/(R_SF_ + R_SIM_). This analysis confirmed that among all groups of neurons, DCP neurons showed the highest selectivity towards SF trials (Figure 5E). Together, these results demonstrate that a subset of A1 neurons was sensitive to SF induced errors.

### Frequency specific error responses are related to the spectral tuning properties in a subset of SF sensitive neurons

As A1 neurons typically respond only to a selective set of frequencies, the error responses induced by SFs could reflect the neuron’s frequency selectivity. We thus investigated if the frequency dependency of the error responses induced by SFs was related to the tuning properties of DCP neurons. Given that we used four different SFs (SF1-SF4), we separated DCP neurons into subgroups based on the particular SF that neurons were most responsive to (Figure 5F). Neurons responded primarily to one SF, which we defined as their best SF. Neurons also showed corresponding, albeit weaker responses, to the same SF in the playback trials, further suggesting that these SF responses could be related to their tuning properties. Out of the total DCP neurons identified responding to SF (n=644, both layers included), 178 showed significant responses to pure tones in the passive setting allowing us to probe the relationship between their tuning properties and their best SFs. Plotting the best SFs against the BFs in these subset of tone responsive neurons showed a positive relationship between best SFs and BFs within the spectral boundary (10 to 40 kHz, Figure 6G). This relationship held both in L4 and L2/3. In addition, we quantified the distance between best SF and the closest frequency that evoked significant responses within the tuning curves of these neurons. The distribution of the distance centered around zero for both L4 and L2/3 (Figure 6G, right). These results suggest that in tone responsive DCP neurons, their SF selectivity is closely related to their tuning selectivity.

As the majority of these DCP neurons were unresponsive to tones in the passive setting, we investigated whether their SF selectivity could be predicted from the tuning properties of the local neuronal population. For this purpose, we defined local BF as the frequency that most neurons responded to within a radius of 100 μm. Plotting the best SF against local BF did not show any obvious relationship between the two measures (Figure 6H) indicating that BSF is independent of local BF. These results show that SF selectivity is best predicted by the neuron’s own tuning while the local population’s frequency selectivity failed to predict SF selectivity, possibly due to the large variability in the local tuning preference (Bandyopadhyay *et al*., 2010; Rothschild *et al*., 2010; Winkowski and Kanold, 2013).

Next, we investigated whether the encoding of SF induced error responses was layer dependent. The fraction of SF responsive DCP neurons (Figure 6I) and the strength of the error responses (as computed by the response bias index between SF and SIM trials, Figure 6E) (Figure 6J) were similar in L4 and L2/3. These results suggest that neurons in L4 and L2/3 encode the SF induced error responses to a similar degree, mirroring the results from DS perturbations (Figure 5I).

### DCP neurons in A1 form functionally connected ensembles

A1 neurons form functionally connected subnetworks (Francis et al., 2022; Francis *et al*., 2018). The introduction of multiple SFs allowed us to investigate the if neurons with different feature sensitivity formed functionally connected ensembles. We thus computed the noise correlation (NC), a measure that quantifies stimulus independent variability among neuronal pairs, either between DCP neurons and neurons sensitive to other features (F, FM and W, cross-group) or within DCP neurons (intra-group). We found that the intra-group NCs among DCP neurons were the highest, while for cross-group NCs, DCP neurons showed higher correlations with W neurons than with F or FM neurons (Figure 6K). We also found a consistently higher NC among neuronal pairs in L4 than in L2/3. These results suggest that DCP neurons form stronger functional networks within themselves while sharing stronger functional connectivity with action sensitive neurons than sensory information sensitive neurons. These functional connections could underlie their highly nonlinear selectivity towards SF perturbations. Together, similar to DS induced DCP neurons, SF induced DCP neurons also encode the deviation of the actual sound from the predicted sound, and thus their responses can be interpreted as the difference between predicted sound generated by the forward model and the actual sensory input (Figure 6L).

### Feature sensitive neurons form spatial clusters

The frequency selectivity among neurons in A1 is heterogenous (Bandyopadhyay *et al*., 2010; Rothschild *et al*., 2010; Winkowski and Kanold, 2013). We thus investigated the spatial distribution of different feature sensitive groups of neurons is also heterogeneous. Figure 7A shows two example FOVs from L4 and L2/3 respectively with identified feature sensitive neurons color coded. To investigate if functionally similar neurons were spatially clustered, we compared the homogeneity index (Deneux *et al*., 2016), which quantifies the fraction of neurons belonging to the same group within a local population, against randomly shuffled data. This analysis revealed that all feature sensitive neurons were more likely to cluster with neurons of the same identity and thus were non-uniformly distributed in both L4 and L2/3 (Figure 7B, C).

**Figure 7.**
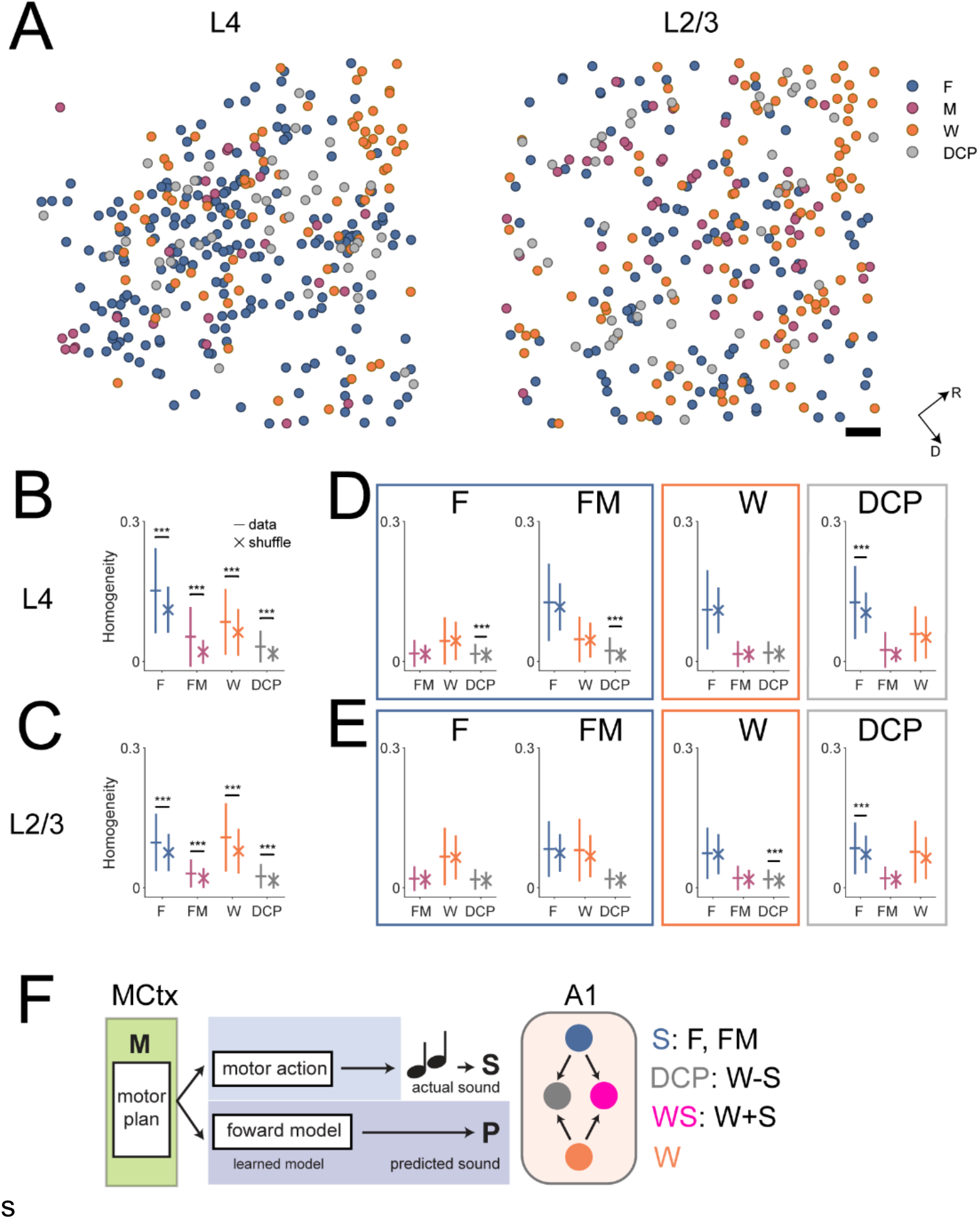
Feature sensitive neurons form local clusters. (A) Two example FOVs from L4 and L2/3 respectively showing the spatial distribution of feature sensitive neurons. DCP neurons contain all identified DCP sensitive neurons, i.e., spectral boundary, DS or SF sensitive neurons. F neurons contain both F and FT sensitive neurons from Figure 2. FM neurons contain both FM and FMT sensitive neurons from Figure 2. W neurons contain both W and WS (excluding boundary sensitive neurons) from Figure 2. The black scale bar represents 100 μm. (B) and (C) The homogeneity indices that quantifies the fraction of local population of neurons that belong to the same category are shown as a function of cortical layer. Data from all FOVs were pooled together. In both L4 and L2/3, each feature sensitive group showed significant within group clustering as actual data showed higher homogeneity indices than shuffled data (Wilcoxon rank sum test, data vs. shuffle, L4: F, p=2.2×10^−128^; FM, p=4.9×10^−29^; W, p=8.2×10^−22^; DCP, 9.4×10^−17^; L2/3: F, p=7.9×10^−41^; FM, p=9.4×10^−10^; W, p=6.3×10^−50^; DCP, 8.9×10^−12^). The markers show the mean while the vertical lines show the 95% confidence interval of the mean. (D) The cross-group homogeneity indices are shown as a function of the feature groups for L4. Each panel shows whether other feature groups spatially cluster in the vicinity of one particular feature group, e.g., the first panel shows the fraction of each group around F neurons. One tailed Wilcoxon rank sum test (significance adjusted with Bonferroni correction), F neurons: F vs. FM, p=1; F vs. W, p=1; F vs DCP, p=4.1×10^−5^; FM neurons: FM vs. F, p=0.21; FM vs. W, p=0.96; FM vs. DCP, p=4.1×10^−5^; W neurons: W vs. F, p=1; W vs. FM, p=1; W vs. DCP, p=0.56; DCP neurons: DCP vs. F, p=2.0×10^−5^; DCP vs. FM, p=0.0026; DCP vs. W, p=0.29. (E) The same as in (D) but for L2/3 neurons. One tailed Wilcoxon rank sum test (significance adjusted with Bonferroni correction), F neurons: F vs. FM, p=0.81; F vs. W, p=1; F vs DCP, p=0.022; FM neurons: FM vs. F, p=0.12; FM vs. W, p=0.028; FM vs. DCP, p=0.061; W neurons: W vs. F, p=0.97; W vs. FM, p=0.081; W vs. DCP, p=5.5×10^−6^; DCP neurons: DCP vs. F, p=8.2×10^−5^; DCP vs. FM, p=0.30; DCP vs. W, p=0.022. (F) The various sound-action interactions could be explained using the framework of forward model, which encodes the predicted sensory outcome (predicted sound, P) from motor corollary signals. The output of such circuits (W neurons) interacts with sensory information encoding signals (F and FM neurons), which could give rise to facilitative responses in DCP-insensitive WS neurons (W+S) as well as deviation responses evoked by the mismatch between sound and action in DCP neurons (W-S).

Given that we identified neurons of different feature groups, we next asked if neurons of different feature groups co-locate with neurons from other feature groups. For this purpose, we computed the fraction of neurons that belonged to other feature sensitive groups than the neuron at the center of the radius and compared the distribution of values against shuffled data (Figure 7D, E). The majority of the comparisons produced insignificant results, which suggest that neurons of different feature sensitive groups were largely randomly scattered across FOVs, despite their within-group spatial clustering, which might facilitate the interaction between neurons encoding different behaviorally relevant information by superimposing their spatial locations.

## Discussion

We trained mice to perform an interactive auditory task while imaging large populations of neurons in A1. We identified separate groups of neurons sensitive to features such as sound, action, reward, and errors. These neurons exhibited layer-specificity for certain features. For example, while L4 contained more sound-encoding neurons, while L2/3 contained more neurons that responded to the mouse’s actions. In contrast, error sensitive neurons were equally distributed across layers.

The sound evoked responses in sound-encoding neurons were typically more suppressed during active sessions than during passive sessions consistent with prior results (Kuchibhotla *et al*., 2017; Otazu *et al*., 2009).

We identified two groups of action sensitive neurons. One group was activated by actions both within trials and during intertrial intervals. However, their responses were more suppressed during trials where sound was presented. The second group of neurons were preferentially driven by the joint presence of action and sound, and we found that a subset of these neurons encodes the spectral boundary of our interactive task. Perturbations of the coupling between action and sound evoked error responses in subsets of A1 neurons. Lastly, feature sensitive neurons were spatially clustered indicating that there is a local microorganization of A1 with respect to the integration of sound and action. Together, our results show that in our interactive behavioral task, a significant portion of A1 neurons were dedicated to encoding action related information, which could give rise to the highly nonlinear responses we observed either at the spectral boundary or evoked by the DS and SF perturbations.

The nonlinear responses of WS and DCP neurons could be unified under the framework of forward model in the motor control theory, i.e., rather than relying on instantaneous sensory feedback for motor control, the brain predicts the sensory outcome by learning a forward model that translates the motor corollary signals into the sensory domain and compares those signals with the incoming sensory input (Kawato, 1999). Under this framework, the responses of WS and DCP neurons could be interpreted as the interactions between the output of the forward model and the feedforward sensory input with opposite signs (Figure 7F). While the responses of DCP-insensitive WS neurons (Figure 4J) could be seen as facilitation between the predicted sound and the actual sound, the responses of various DCP neurons (Figure 4J, 5D, 6D) could be seen as suppressive between the predicted and actual sound (W-S). Indeed, the presence of sound encoding neurons (F, FM) and action encoding neurons (W) provides a substrate for the forward model and downstream nonlinear interactions in WS and DCP neurons. Therefore, our results are consistent with the presence of a learnt forward model within A1 that actively predicts the sensory outcome of ongoing action of the animal.

Our interactive behavioral design allowed us to manipulate the coupling between action and sound feedback, such that error responses could be evoked. This approach is inspired by studies in the visual system where visual flow mismatch evoked error responses in the mouse visual cortex (Keller *et al*., 2012; Zmarz and Keller, 2016). In the auditory system, efforts have been made to couple the animal’s action, e.g., locomotion or lever pushing, with the auditory feedback (Audette et al., 2021; Rummell *et al*., 2016; Schneider *et al*., 2018). In these studies that employed a simple coupling, the sound was often only secondary to the task and might not necessarily require attention to the sound per se (Audette *et al*., 2021). Our task introduced a more complex mapping between sound and action while requiring the animal to pay attention to the sound presented, i.e., the direction of the turn depends on the initial frequency of the sound stream.

In our task, we used wheel turning to report the animal’s behavior. We monitored wheel turning throughout the entire experiment session and together with the sound information, we were able to rule out responses due to sound and action while identifying putative error responses. Consistent with previous reports (Musall et al., 2019), we observed considerable action induced responses in A1 and identified action sensitive neurons through linear regression. Moreover, our large imaging field enabled us to identify a subset of A1 neurons (~1.3% of all imaged neurons) responding specifically to reward consumption and these responses were robust, which is consistent with a previous study reporting licking induced neural activities in a subset A1 neurons (Clayton *et al*., 2021). However, these neurons were rare and not a majority of A1 neurons (Francis *et al*., 2018) and auditory responses also show heterogeneous modulations by motor signals in A1 (Henschke et al., 2021).

We identified A1 neurons with specific selectivity to wheel turning directions that act in essence like specific motor neurons. We speculate that these neurons receive long range input from motor related areas. Both primary (M1) and secondary motor cortex (M2) have projections to A1, and M2 innervate both pyramidal and parvalbumin positive neurons in A1, although the net effect of M2 activation in A1 could be feedforward inhibition (Nelson *et al*., 2013; Schneider et al., 2014). Moreover, the basal forebrain also projects to A1 and these cholinergic inputs in A1 are active during movement (Nelson and Mooney, 2016). Given the positive sign of the action responses, it is thus possible that the action sensitive neurons receive prominent input from basal forebrain. Action neurons are among the least tone responsive, and the selectivity of the turn directions suggests that action neurons were not activated by the sound caused by the movement. Thus, these action neurons seem to provide a functional motor network within A1 that broadcasts action specific signals to other A1 neurons, potentially giving rise to the action dependent error responses.

We identified both action driven and sound driven error responses in A1. The action driven error responses were evoked when the carrier frequency change failed to follow the wheel turning activities, which happened either due to the boundary of the frequency spectrum or due to the introduction of the perturbations. The error responses evoked by these decoupling events thus represent a highly nonlinear sensitivity to the combination of a stationary carrier frequency and concurrent wheel turning. These error responses could be interpreted as signaling the violation of the fundamental rule of the task, i.e., turning the wheel changes the carrier frequency. On the mechanistic level, these responses could result from the nonlinear interactions between action encoding and sound encoding neurons. In DS trials, the action driven error responses were time locked to the onset of the action, while in SF trials, the error responses occurred after the SF onset. These phenomena further suggest that these responses are due to low-level sound or action selectivity, but such selectivity was highly dynamic and dependent on previous action and sensory history.

One of the key hypotheses of the predictive coding theory is that cortical layers could form hierarchical structures in terms of error signaling (Heilbron and Chait, 2017). Despite the fact that more L2/3 neurons were action sensitive, we did not find a systematic shift in the error response strength across L4 and L2/3. This finding suggests that cortical layers might not be the minimum functional unit in terms of generating error responses, and it is possible that the proposed hierarchy exists only across different cortical regions.

In conclusion, we designed an interactive auditory task that allowed us to probe complex interactions between sound and action in mouse A1. We identified both sensory information and action sensitive neurons in A1, as well as neurons signaling reward and error responses evoked by the decoupling between action and sound. Our approach potentially provides a new approach in terms of studying predictive coding in the auditory system.

## Acknowledgements

We thank Dr. Clare Jung Yoon Choi for the useful discussions on the project and Dr. Shihab Shamma for comments on the manuscript. Supported by U19 NS107464 and R01DC17785.

## Methods

### Animals

All protocols and procedures are approved by the Johns Hopkins Institutional Animal Care and Use Committee. In this study we trained and imaged 3 male and 3 female adult mice which were F1 generation of Thy1-GCaMP6s (JAX# 024275) crossed with CBA/CaJ mice (JAX# 000654). The F1 offspring has both widespread expression of GCaMP6s in excitatory cells and minimal hearing loss throughout their lifespan (Frisina et al., 2011; Shilling-Scrivo et al., 2021). The mice used in this study ranged from 4 to 8 months old. All mice were reared in a 12-h light/12-h dark cycle in the institutional animal colony. Food was provided *ad libitum* while water intake was restricted for behavioral training.

### Behavior Paradigm

The mice were trained on an interactive behavior paradigm. Specifically, the mice were presented with a frequency stream which started with either 10 kHz (low starting frequency) or 40 kHz (high starting frequency). By turning the wheel (63 mm in diameter) placed beneath their front paw, mice were able to modulate the carrier frequency by either turning to their right (carrier frequency increases) or turning to their left (carrier frequency decreases). The lower and upper bound of the carrier frequency was defined by the low and high starting frequency, respectively, beyond which left turning at lower frequency bound or right turning at upper frequency bound) produced no carrier frequency change. If in a given trial the mouse was presented with the low starting frequency (10 kHz), a hit was achieved if the mouse turned the wheel to its right and the carrier frequency entered a high frequency target zone (0.25 octave wide, between 33.6 to 40 kHz) and remained in the target zone for at least 0.2 second. For trials with a high starting frequency (40 kHz), the target zone was located at the low frequency end (10 to 11.9 kHz) and thus, the mouse was required to turn left. The animal was given a maximum of 6 seconds to complete each trial. 22.5 degrees of rotation corresponded to 1 octave change in carrier frequency. After a hit was achieved, the frequency stream continued for 0.5 second while its carrier frequency remained the same as the time of hit. During this period, a 10 Hz amplitude modulation was added to the frequency stream to reinforce the stimulus salience with the behavioral relevance. After the sound terminated, a servo-controlled waterspout was elevated, and one drop of water (~5 μl) was dispensed. The mouse had 2 seconds for reward consumption before the waterspout was retracted by the servo. If the mouse turned beyond 30 degrees in the incorrect direction, the trial was deemed incorrect and terminated. If the mouse failed to reach the target zone without turning incorrectly (beyond 30 degrees in the incorrect direction), the trial was deemed a miss. Both incorrect and miss trials were punished with an 8-second timeout. There was a fixed 0.5 second period between the end of reward or punishment period of the previous trial and the start of the next trial. At the beginning of each trial, the mouse was monitored for any spontaneous wheel turning activity and only after the mouse remained inactive for another 0.5 second was the sound presented. The program waited indefinitely until this criterion was met. After the sound started, the first 0.1 second was considered a ‘grace period’ where any wheel turning was discounted, and thus wheel turning during this period did not produce carrier frequency change. The behavior program was implemented with LABVIEW 18.0 and was run on NI-USB 6343 (National Instrument).

During imaging, we controlled the onset of any sound such that it roughly aligned with the frame start trigger from the 2P imaging system. The jitter between the onset of the sound and the onset of one particular frame was around 0.17 ± 1.23 millisecond. This approach allowed us to more precisely align neuronal responses with the animal’s behavior. Furthermore, we employed a second device (NI PCI-6251) that used the same 2P frame start triggers as the acquisition clock to record a copy of the rotary activity or wheel movement. Thus, not only did we record the wheel movement within each trial at 100 Hz but we also recorded the wheel movement at 15 Hz (the 2P frame rate) throughout the entire imaging session.

### Perturbation trials during behavior

To test the hypothesis that A1 encodes error signals induced by mismatch between the behavior and the sensory outcome, we introduced two forms of perturbations: Stop-Frequency (SF) and Delay-Sound (DS). In SF perturbations, we chose 4 frequencies outside the target zone (13.2, 17.4, 23.0, 30.3 kHz). These frequencies were symmetrical about the center of the spectral range (20 kHz) in the logarithmical space and equally spaced by 0.4 octave. During the SF perturbation trials, as the frequency trajectory reached one of the SFs, the carrier frequency was frozen at that SF for 0.25 second, artificially introducing a decoupling period. At the end of this period, the paradigm resumed the normal coupling between the wheel movement and the carrier frequency. We presented SFs in blocks. Every 4 SFs form a block and within each block all 4 different SFs were presented. The order of the 4 different SFs within a block were randomized. We also balanced the starting frequencies by flipping them every time the same SF were encountered, i.e., if one particular SF was introduced in a low starting frequency trial in the previous block, it would be introduced in a high starting frequency trial in the current block and vice versa. We required that between SF trials, there would be a minimum of 2 hit trials and a maximum of 7 hit trials. Within those limits, it was determined with a probability of 0.5 whether SF would be introduced in each trial. However, if the animal failed to reach the SF in a given SF trial, in which case the perturbation effectively did not happen, the SF was carried over to the next trial with the same starting frequency. In all our imaging sessions with SF perturbations, we had a total of 31.4 ± 4.9 (mean±std) SF trials per session, which corresponded to 15.9% ± 1.4% of all trials.

In a different set of imaging sessions, we introduced Delay-Sound (DS) perturbations. In such trials, we introduced a delay of 1 second between the action and the sensory feedback, i.e., the carrier frequency change corresponding to any wheel movement was delayed by 1 second. To balance trials with low and high starting frequencies, every DS trial had the opposite starting frequency of the previous DS trial. Similar to SF trials, we introduced DS with a probability of 0.5 if there were at minimum 3 hit trials but no more than 8 hit trials since the last DS trial. However, if the outcome of a particular DS trial was miss (the animal did not turn sufficiently in the correct direction) or incorrect (the animal turned in the wrong direction and effectively produced no carrier frequency change), we deemed these trials as insufficiently perturbated and repeated the DS trial until the animal turned in the correct direction with a sufficient amount, which typically resulted in a hit. In our imaging sessions with DS perturbations, we had 21.2 ± 6.5 (mean±std) DS trials, which corresponded to 11.6% ± 1.9% of all trials. Both SF and DS perturbations were only introduced during imaging sessions.

### Behavioral training

Mice first received headplate implantations and were allowed 5-7 days of recovery before the start of water deprivation. We restricted the water intake of mice to no more than 1 ml per day. The weight of mice dropped steadily within around 7 days to about 80% to 85% of the original weight. In the meantime, mice were introduced to the behavioral chamber and were accustomed to head-fixation before the training began. We devised 4 stages for mice to systematically advance. The parameters of these stages were overall similar to those of the imaging sessions. The first 3 stages differed in the gain of the wheel, with each stage requiring the mouse to turn twice as much as the previous stage. Specifically, the gains were 5.625, 11.25, and 22.5 degrees per octave change in carrier frequency. In these stages, mice were rewarded as soon as a hit was achieved, and thus they consumed the reward during the period when the amplitude modulation was added to the frequency stream. In the 4^th^ stage, the reward time was delayed to 0.5 second after the end of the frequency stream in a hit trial and thus separated the reward consumption behavior from sensory input in time. The rotary gain of the 4^th^ stage was the same as in the 3^rd^ stage. The mice advanced to the next stage if they completed at least 100 trials within a session and if the maximum hit rate over any consecutive 100 trials were above 60%. This relatively relaxed criterion was chosen such that mice were not overtrained on intermediate stages and could advance to the final stage more quickly, which shared the parameters of the actual experimental session. After mice reached a performance of about 70% to 80%, they were implanted with cranial windows and transferred to a behavioral setup under 2-photon imaging.

### Sound stimulus

During the task, the carrier frequency of the sound stream increased or decreased dynamically as the mouse turned the wheel towards its right or left, respectively. The carrier frequency was bounded between 10 and 40 kHz. The wheel rotation was detected via a rotary encoder and the carrier frequency was updated 100 times per second. The carrier frequency changed with a step of 1/128 octave. Every 10 ms, the program translated the change in rotary reading (ΔR) into the change in the carrier frequency (ΔCF) and output a linear frequency sweep of 10-ms duration with its start at the current carrier frequency (CF) and its end at the CF+ΔCF. Therefore, the sound stream was essentially a piecewise linear frequency sweep that updated every 10 ms. At each time point, the rotary reading was smoothed over the previous 100 ms with a mean filter to produce a smoother frequency stream. The amplitude of the frequency stream at any time point was interpolated using an array of frequencies and their calibrated amplitude at 60 dB SPL. Within the frequency bounds, 22.5 degrees of rotation mapped onto 1 octave change in carrier frequency.

Right after the conclusion of the active session, we presented the mice with pure tones that ranged from 7.1 to 56.5 kHz with 0.25 octave spacing at the same sound level as in the active session. These tones were used to construct tuning curves under the passive condition. After the presentation of the pure tones, we presented mice with the playback of the frequency streams selected from a subset of trials in the previous active session. These playback sound streams were generated by reading the stored rotary positions as if they were generated in realtime by the animal. Specifically, all perturbation trials were selected along with 10 hit trials, 6 incorrect trials and 6 miss trials which were evenly split between trials with low or high starting frequency. These selected trials were repeated 4 times in a random order.

The sound waveform was generated by NI 6343 (National Instrument), which was used as input for ED1 speaker driver (Tucker-Davis Technologies) that drove an ES1 open field speaker (Tucker-Davis Technologies).

### Widefield imaging

To locate A1, we performed widefield imaging as previously described (Liu et al., 2019). In short, we used a blue LED of 470 nm wavelength (M470L3, Thorlabs Inc.) to illuminate the cranial window while imaging the excitation light with a PCO Edge 4.2 camera. We presented 5 tones ranging from 4 kHz to 64 kHz with 1 octave spacing and at 3 sound levels (30, 50 and 70 dB SPL). The images were of size 330 by 330 in pixel and were sampled at 30 Hz. We identified A1 by identifying the low to high frequency gradient starting from the caudal side of the cranial window towards the rostromedial side.

### Two-photon imaging and analysis

We performed two photon imaging in A1 while the mouse performed the task under the microscope (Bruker Ultima 2Pplus). We imaged A1 with a 16X Nikon objective (NA 0.80) and at an optical zoom of 1X. The field of view was of size 1109.9 by 1109.9 μm. The frame rate was 15 Hz. During the experiment, the head of the mouse was upright while the microscope nosepiece was rotated from the vertical position by about 50 degrees to match the angle of the cranial window surface. The imaging laser (Coherent Discovery HP) was tuned to 920 nm wavelength.

For analysis, we used Suite2P package to perform motion correction, automated regions of interest (ROI) detection and raw cellular and neuropil fluorescence trace extraction (Pachitariu et al., 2016). We corrected the neuropil contamination using the following equation:

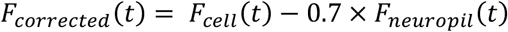

To convert the time-varying neuropil-corrected fluorescence trace into ΔF/F traces, we computed the baseline using the same method as described before (Liu and Kanold, 2021). In short, as most excitatory neurons had sparse firing, the fluorescence would fluctuate around the baseline during most of the imaging session. Thus, for each neuron, we constructed the histogram of fluorescence values over time and identified the value that appeared most often at the resolution of the histogram binning. We used this value for that neuron’s baseline and then computed ΔF/F over time using the following equation:

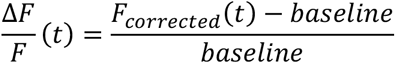

For a subset of our analyses, we used inferred spikes from the ΔF/F traces. The spikes were extracted using OASIS package (Friedrich et al., 2017).

### Cranial Window Surgery

We followed a similar procedure for cranial window implant as previously described (Liu and Kanold, 2021). In short, 0.1 ml dexamethasone (2mg/ml, VetOne) was injected subcutaneously 2-3 hours before the surgery started in order to prevent brain swelling during surgery. All surgery tools were sterilized using a bead sterilizer (18000-45, Fine Science Tools). We anesthetized mice with isoflurane (Fluriso, VetOne) using a calibrated vaporizer (Matrix VIP 3000). We used 4% for induction and 1.5-2% for maintenance. During surgery the body temperature of the animal was maintained at 36.0 degrees Celsius. Next, we exposed the bone covering the auditory cortex by removing the skin and the muscles. A circular craniotomy was then performed over the left auditory cortex with a diameter of ~3.5mm using a dental drill. We then placed a custom-made cranial window over the exposed brain. The window consisted of 2 layers of 3mm round coverslips (64-0720, CS-3R, Warner Instruments) stacked at the center of a 4mm round coverslip (64-0724, CS-4R, Warner Instruments) and secured with optic glue (NOA71, Norland Products). The edge of the cranial window was then sealed with Kwik-sil (World Precision Instruments). Finally, C&B-Metabond (Parkell Inc.) was applied to secure the window. After the surgery, 0.05 ml Cefazolin (1 gram/vial, West Ward Pharmaceuticals) and 0.1 to 0.15 ml Carprofen (1 mg/ml) was injected subcutaneously, and the animal recovered under a heat lamp for 30 minutes before being returned to the home cage. For each of the two days after the surgery, the animal was given additional 0.05 ml dexamethasone (2mg/ml) and 0.1 to 0.15 ml Carprofen (1 mg/ml). The animal was taken off training schedule for 3 to 5 days after the surgery and given supplemental water.

### Linear model construction and fitting

We constructed linear models that included several factors to explain the neuronal responses. In our first model (Figure 2A), the predictors were constructed from both trials and inter-trial periods. Each row of predictors was constructed from a 0.5 second window, and each window was shifted by 0.25 second. The predictors include frequency energy in different frequency bins (F), the frequency modulation (FM) rate (FM) and wheel movement (W). The predictor F could be thought of as a spectrogram with a low spectral resolution. Specifically, the boundaries of the 5 bins we used were 0, 0.25, 0.625, 1,1.375, 1.75, 2, which were measured in octaves relative to the 10 kHz, the lower bound of the carrier frequency. The first ([0, 0.25]) and last bin ([1.75, 2]) corresponded to the low and high frequency target zone, respectively. The rest of the 4 bins were of equal size: 0.375 octave. For each of the time window the predictors were extracted from, we quantified the proportion of time the carrier frequency resided in each of the 6 bins, in order to proximate the energy in each frequency bin. Next, we calculated the S predictor first by taking the derivative of the frequency trajectory represented in octave. We then constructed two histograms for positive and negative FM rate respectively. For positive FM rate, we used the following edges for the histogram: 0, 5, 10, 20 oct/sec. For negative FM rate, we used the same edges with the opposite sign. In both cases, 0 oct/sec was not included in the histogram and thus only strictly non-zero FM rate was considered. For each time window, such histograms were constructed, which measured the occurrence of different FM rate. For the W predictors, we took the first order derivative of the rotary traces and summed positive and negative terms separately, which produced +W and -W, representing the absolute size of right and left turning events as measured in degrees. To model the effect of behavioral state, we introduced one “Task” term (T) that took value of 1 for predictor values constructed from active trials while taking value of −1 for predictor values constructed from playback trials. We then included the cross-product terms between sound encoding terms, i.e., F and FM, and T to produce FT and MT predictors. If a particular neuron responded preferentially to F during active but not playback trials, then one would expect positive F coefficients and positive FT coefficient. Such interaction term was not constructed for W term as the animal showed little activity during the playback session. Next, to model the interaction between sound and action, we introduced the WS term, which was the W terms (separated according to left and right terms) times the summation of all frequency energies (summing over all F terms). Thus, the WS term was equivalent to the W terms windowed by the presence of sound. Finally, we also included a reward term (Rw) that marked the presence of reward. For this model, the design matrix had 6 F terms, 6 FT terms, 6 FM terms, 6 FMT terms, 2 W terms, 2 WS terms, 1 Rw term and 1 T term. Each active and playback session pair would produce ~10,000 observations for the row of the design matrix. We used inferred spikes as the neuronal responses for the model due to the more accurate timing. Next, we fit our linear model using LASSO (MATLAB built-in function, lasso), which had the benefit of automated feature selection. For each neuron, we used a 10-fold cross-validation to determine the optimal regularization term value that achieved the minimum error, and subsequently used the same value for fitting all the observations.

To determine the sensitivity of neuronal responses to the different predictors, we calculated the ΔR^2^s by measuring the difference between full model R^2^ and the R^2^ calculated from shuffling a particular set of predictor values. For example, to determine the ΔR^2^ of the F predictor, all 6 F terms were shuffled. Next, we grouped neurons based on which predictor had the highest ΔR^2^. We also required that the maximum ΔR^2^ cross a threshold of 0.05.

For Delay-Sound (DS) perturbations, we extracted predictors from 4 non-overlapping time-windows of 0.5-second duration (Figure 5A). The first window was aligned with the onset of the action. Due to the 1-second delay between action and carrier frequency change, in the first second following action onset, the carrier frequency remained the same as the starting frequency. Thus, the first two time-windows correspond to this period, which we define as the action window, while the other two time-windows corresponded to the period when the delayed carrier frequency change occurred. We define this window as the sound window. Next, we extracted action driven decoupling (DCP) in the action window and sound driven DCP in the sound window. Action driven DCP (DCP_A_) was defined as the period of time when the wheel turning failed to translate into carrier frequency change. Sound driven DCP (DCP_S_) was defined as the period of time when the frequency change did not correspond with the concurrent action (Figure 5B). We broke down DCP_A_ by the starting frequency at which it occurred (low vs. high) and we broke down DCP_S_ by the direction of the frequency sweep (up vs. down). All other predictors were constructed as in the first model. For the rows of the design matrix, we included observations from DS trials and the corresponding playback trials. Only DS trials with a delayed hit were included. We also included non-DS (NDS) trials where hits were achieved. These trials with hits were selected such that the responses due to the action could be separated from the genuine error responses. In total, the design matrix had 31 columns (6 F terms, 6 FT terms, 6 FM terms, 6 MT terms, 2 W terms, 2 DCP_A_ terms, 2 DCP_S_ terms and 1 T term) and ~800 rows.

For our third model that focused on the Stop-Frequency (SF) perturbation, we used the same approach and constructed similar predictors. As the duration of the SF perturbation was 0.25 second, we constructed our predictors from two time-windows of 0.25 second. The first window spanned the 0.25 second before SF onset, while the second window spanned the 0.25 second after the SF onset. All predictors but DCP were constructed in the same fashion as in the first model. As SFs happened outside target zones, we computed the action driven DCP in the 4 middle frequency bins that we used for F predictors, where the 4 SFs were within each of the 4 bins. Thus, this model had 4 DCP terms, one for each of the 4 SFs. In total, the design matrix for the SF perturbations had 31 columns (12 F and FT terms, 12 FM and MT terms, 2 W terms, 4 DCP terms, 1 T term). For rows in the design matrix, all SF perturbation trials in the active sessions were included. A typical active session included ~30 SF trials. We next included the playback trials that corresponded to SF trials in the active sessions. As each SF trial was played back with 4 repeats, the playback sessions added ~120 observations to the design matrix. To further account for the contribution of pre-SF period to neuronal responses, we included so-called SIM trials, where the frequency trajectory was similar to that of the SF trial immediately before SF onset (Figure 6B). We hypothesize that if the responses during the post-SF period were due to pre-SF factors, then the responses to SF trials would be similar to those in SIM trials. For each of the 8 unique SF trials (4 SFs × 2 starting frequencies), we each selected the closest n matching SIM trials, such that the total number of SIM trials (8n) were 5 times that of SF trials (~150 observations). In the end the design matrix had about 300 rows.

In Figure 4H, in order to determine the feature to which the neural responses showed a higher temporal sensitivity, we constructed the predictors with the same temporal resolution as the neural responses. First, we expressed the wheel turning events and DCP events by shrinking these events to their onset time with the corresponding event size (Figure 4H, left top). Next, we convolved these events with a set of 9 B-spline basis spanning 2 seconds (Figure 4H, left bottom) to get the predictors (Figure 4H, right). Such constructed predictors could be used to capture the temporal profile of the neural responses.

### Local Best Frequency

In Figure 5I, we investigate whether BSF depended on the Local Best Frequency (LBF), which measured the tuning preference of local population of neurons. To compute LBF, we first identified the neurons within a 100-μm radius of the neuron in question. Next, we summed the tuning curves across individual neurons weighted by the respective significance, i.e., significant responses had a weight of 1 while non-significant responses had a weight of zero. Thus, we obtained a population tuning curve whose peak represented the frequency that evoked the most responses in the local group of neurons, which we defined as LBF. The significance of tone evoked responses for each neuron was determined similar to described before (Liu and Kanold, 2021). In short, we constructed the 99.9% confidence interval of the pre-stimulus-onset and post-stimulus-onset ΔF/F values, and one tone is considered significant if the corresponding confidence intervals were non-overlapping. For pre-stimulus-onset period, we chose a window of ~0.5 second duration immediately before the tone onset. For post-stimulus-onset period, we chose a window of ~0.5 second duration offset by ~0.27 second (4 frames) from the tone onset, in order to better capture the peak of the ΔF/F trace.

### Spatial Clustering Analysis

In Figure 7, we quantified the spatial clustering of different feature sensitive group following a similar strategy in Deneux et al. (2016). We calculated a homogeneity index for each neuron, which quantified the proportion of neurons within a 100 μm radius that belonged to either the same group as the neuron at the center or belonged to the other feature sensitive groups. We determined the significance of the distribution of the homogeneity indices by comparing the actual distribution against the shuffled data, where we randomly assigned the feature sensitive identity to neurons in the FOV. The values of real and shuffled data were pooled from all FOVs and compared to determine significance.

