## Supplemental Figures for "Interactive auditory task reveals complex sensory-action integration in mouse primary auditory cortex"

**Supplementary Figures 1-3**

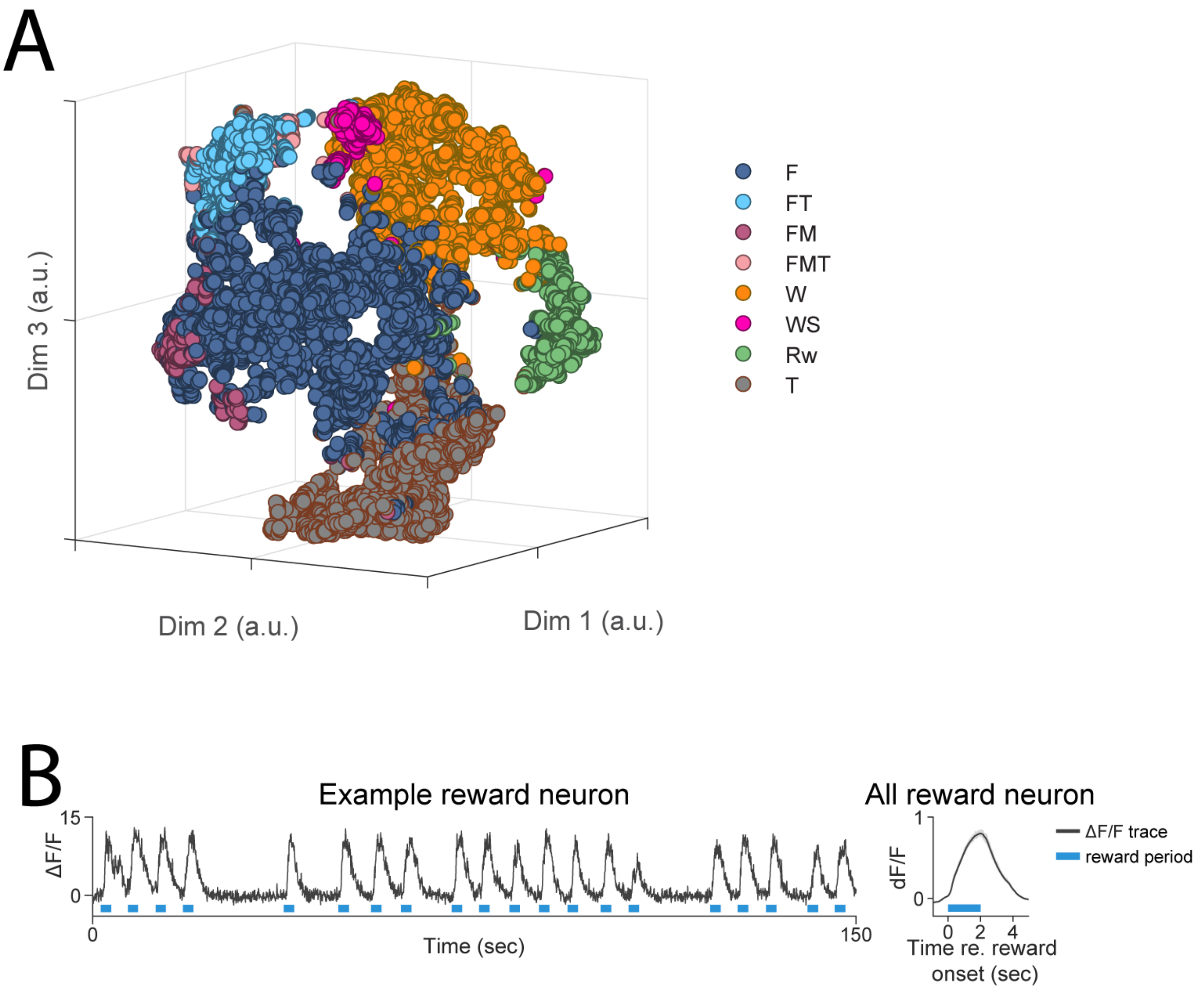


**Supplemental Figure 1 (Related to Figure 2)**

(A) 3D scatter plot showing the identified feature sensitive groups of neurons projected on the lower dimensional space generated by the T-sne algorithm. (B) Left: the fluorescence trace from an example reward sensitive neuron. The blue lines mark the reward consumption period. Right: the traces from all reward neurons aligned to the onset of the reward period. The solid line represents the mean and the shaded region represent the 95% confidence interval constructed through bootstrapping.

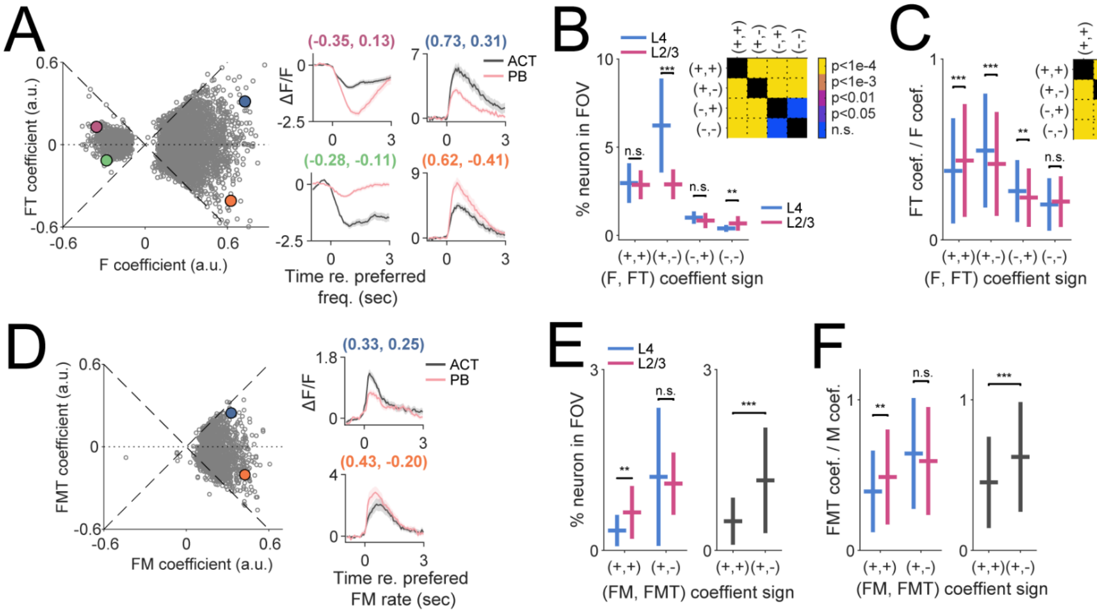


**Supplemental Figure 2 (Related to Figure 2)**

(A) Left: scatter plot the coefficients of the F and FT term from both F and FT group. The four quadrants of neurons showed different levels of behavioral state dependence in their frequency responses. For example, while positive F terms indicate general excitatory neuronal responses driven by the preferred frequency channel (blue and orange scatter points show two such example neurons), a positive FT term indicates stronger responses during the active (ACT) session (blue scatter example) and a negative FT term indicates stronger responses during the passive playback (PB) session (orange scatter example). The purple and green scatter points represent two neurons with negative F terms, indicating that they responded to the preferred frequency with a decrease in fluorescence. The dashed lines were of slope 1 and -1, respectively. Right: the traces from four example neurons corresponding to the 4 colored scatter points on the left are shown. The colors of the titles match the four colored scatter points shown on the left. The traces are aligned to the onset of their preferred frequencies and vertically offset by their immediate pre-onset baselines. (B) The fraction of F, FT neurons as a function of both the sign of the (F, FT) coefficient pair and cortical layer. L4 vs L2/3 Wilcoxon rank sum test:

(+,+), $p=0.77$; (+,-), $p=9.2\times10^{-8}$; (-,+), $p=0.066$; (-,-), $p=0.0058$. The inset shows the multiple comparison between neurons with different (F, FT) coefficients signs regardless of the cortical layer. The (+,-) group had the most number of neurons. (C) The ratios between the absolute values of FT and F coefficients are plotted as a function of both the sign of the (F, FT) coefficient pair and cortical layer. L4 vs L2/3 Wilcoxon rank sum test, (+,+), $p=7.6\times10^{-7}$; (+,-), $p=3.5\times10^{-13}$; (-,+), $p=0.012$; (-,-), $p=0.12$. The inset shows the multiple comparison between neurons with different (F, FT) coefficients signs regardless of the cortical layer. The (+,-) group showed the strongest modulation between ACT and PB session. (D) The same as in (A) but for FM and FMT group. Unlike F and FT group, the FM coefficients were predominantly positive. (E) The fraction of FM, FMT neurons as a function of both the sign of the (FM, FMT) coefficient pair and cortical layer. L4 vs L2/3 Wilcoxon rank sum test: (+,+), $p=0.0073$; (+,-), $p=0.75$. Right: comparison between (+,+) and (+,-) group regardless of the cortical layer, Wilcoxon rank sum test, $p=1.2\times10^{-9}$. (F) The ratios between the absolute values of FMT and FM coefficients are plotted as a function of both the sign of the (FM, FMT) coefficient pair and cortical layer. L4 vs L2/3 Wilcoxon rank sum test: (+,+), $p=0.0023$; (+,-), $p=0.0285$. Right: comparison between (+,+) and (+,-) group regardless of the cortical layer, Wilcoxon rank sum test, $p=9.8\times10^{-18}$.

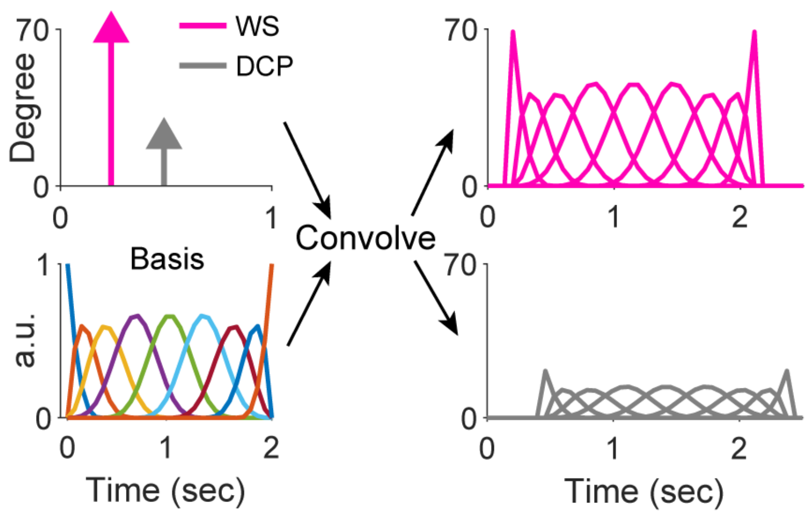


**Supplemental Figure 3 (Related to Figure 4)**

Left top: we expressed the wheel movement during sound (WS) and DCP from the same example as in (Figure 4A) by assigning the amplitude of these events (measured in degrees) to the respective onset time. In this example, the left turning event was of ~70 degrees from the motion start to end. The DCP event was of size ~20 degrees, i.e., ~20 degrees of turning happened on the spectral boundary and thus were not translated into carrier frequency changes. These events were then convolved with a B-spline basis set spanning 2 seconds (left bottom) to construct predictors with the same temporal resolution as the imaging frame rate (right, top and bottom).
